# Fluvoxamine maleate ameliorates Alzheimer disease pathology by mitigating amyloid-beta load and neuroinflammation in 5XFAD mice

**DOI:** 10.1101/2023.12.17.572086

**Authors:** Sukhleen Kaur, Kuhu Sharma, Ankita Sharma, Kamalpreet Kaur Sandha, Syed Mudassir Ali, Riyaz Ahmed, P. Ramajayan, Parvinder Pal Singh, Zabeer Ahmed, Ajay Kumar

## Abstract

**Background:** Alzheimer pathology (AD) is accompanied by the deposition of amyloid beta (Aβ) and chronic neuroinflammation, where NLRP3 inflammasome is particularly involved. In this study, we found that the OCD drug fluvoxamine maleate (FXN) can potently ameliorate AD pathology in 5XFAD mice by autophagy-mediated clearance of Aβ and inhibition of NLRP3 inflammasome.

**Methods:** We used mice primary astrocytes to establish the mechanism of action of FXN against NLRP3 inflammasome by using various techniques like ELISA, Western blotting, confocal microscopy, Immunofluorescence, etc. The validation of the anti-AD activity of FXN was done in transgenic 5XFAD mice after two months of treatment followed by behavior analysis and studying inflammatory and autophagy proteins along with immunohistochemistry analysis for Aβ load in the hippocampi.

**Results:** Our data showed that FXN induces autophagy to inhibit NF-κB and NLRP3 inflammasome at a low concentration of 78 nM apart from directly inhibiting NLRP3 inflammasome in primary astrocytes. FXN activated the PRKAA2 pathway through CAMKK2 signaling, which led to the induction of autophagy in primary astrocytes. FXN inhibited the ATP-mediated NLRP3 inflammasome through autophagic degradation of NF-κB and thus caused the downregulation of pro-IL-1β and NLRP3. The anti-NLRP3 inflammasome effect of FXN was reversed when autophagy was inhibited either by genetic knockdown of the PRKAA2 pathway or by bafilomycin A1.

Furthermore, FXN treatment led to improved AD pathology in 5XFAD mice, which displayed a significant improvement in multiple behavior parameters like working memory and neuromuscular coordination and they behaved more like wild-type animals. We found that FXN improved behavior in 5XFAD mice by clearing the Aβ deposits from the hippocampi along with a significant reduction in multiple inflammatory proteins, including NF-κB, GFAP, IBA1, IL-1β, TNF-α, and IL-6 associated with NF-κB and NLRP3 inflammasome in the brain. Moreover, these changes were accompanied by increased expression of autophagic proteins.

**Conclusion:** Our data suggest that to ameliorate AD pathology, FXN simultaneously targets two key pathological features of AD that is Aβ deposits and neuroinflammation. Being an approved drug, FXN can be pushed as a potential drug candidate for human studies against AD.

## Background

Deposition of amyloid beta (Aβ) and neuroinflammation are essential pathological features of Alzheimer disease (AD). Aβ contributes to neuroinflammation through multiple mechanisms. It activates NLRP3 inflammasome either directly by acting as a DAMP for sensor NLRP3 and indirectly by activating TLR4 mediated induction of NF-ĸB pathway [1–6]. It can also activate NLRP3 inflammasome by inhibiting PRKAA2 and autophagy [7]. Inflammation, in turn, also promotes the Aβ pathology through various pathways. NLRP3-mediated activation of IL-1β inhibits the microglial cells from engulfing the Aβ plaques and thus contributes to its enhanced deposition [8–10]. ASC specks secreted by immune cells directly bind to the Aβ and augment the accumulation of Aβ aggregates [11]. NF-ĸB can also increase the activity of BACE1 to enhance the production of Aβ [12]. Therefore, it’s a kind of vicious cycle which helps in the progression of AD. Many studies have shown that autophagy is a common link between Aβ deposition and activation of NLRP3 inflammasome [13, 14]. Autophagy, by virtue of its ability to recycle the cellular contents, can help clear protein aggregates and limit the activation of NLRP3 inflammasome by reducing the availability of proteins linked with inflammasome. Therefore, autophagy has emerged as one of the most important mechanisms that regulate NLRP3 inflammasome [15, 16]. Some recent studies have shown that induction of autophagy by pharmacological or non-pharmacological means can help mitigate NLRP3 inflammasome mediated or other types of inflammation [17–20]. Therefore, targeting NLRP3 inflammasome and autophagy together can be a good strategy, particularly when most of the monotargeted therapies have not shown any significant clinical advantage for treating AD.

In this study, as a part of our program related to the repurposing of drugs, we identified fluvoxamine (FXN) as a potent inhibitor of NLRP3 inflammasome and NF-ĸB pathway. We show here that the anti-inflammatory properties of FXN are regulated by PRKAA2-mediated autophagy. We attempted to utilize the ability of FXN to simultaneously target inflammation and autophagy in mitigating AD pathology in transgenic 5XFAD mice. We found that FXN can clear Aβ from the hippocampus of mice at a minimal dose 5 mg/kg along with a significant decline in neuroinflammation. These molecular changes in the brain helped improve working memory and overall brain health in 5XFAD mice. Our data projects FXN as a lead drug that should be explored for its possible development for the treatment of Alzheimer disease.

## Methods

### Chemicals and antibodies

Chemicals-RPMI-1640 (Sigma-Aldrich-R6504),Dulbecco’s modified eagle medium (Sigma-Aldrich-D1152), Penicillin (Sigma-Aldrich-P3032), Streptomycin (Sigma-Aldrich-S6501), Sodium bicarbonate (Sigma-Aldrich-S5761), Fetal Bovine Serum (FBS) (GIBCO-10270106), Phosphate buffer saline (PBS) (Sigma-Aldrich-D5652), EDTA (Invitrogen-15575038), Lipopolysaccharide (LPS) (Sigma-Aldrich-L3129), Adenosine 5’-triphosphate (ATP) (Sigma-Aldrich-A6419), Acrylamide (MP Biomedical-193982), Glycine (MP Biomedical-194825),Triton X-100 (Sigma-Aldrich-T8787), Albumin Bovine Fraction V (MP Biomedical-160069), Phenylmethylsulfonyl fluoride (PMSF) (MP Biomedical-195381), Skimmed milk (Himedia-GRM1254), Strataclean resin (Agilent-400714-61), Bafilomycin A1 (Sigma-Aldrich-B1793), Rapamycin (Sigma-Aldrich-R8781),DAPI (Sigma-Aldrich-D9542), glycerol (Sigma-Aldrich-G5516), Tween 20 (Sigma-Aldrich-P7949), HEPES (Sigma-Aldrich-H3375), Paraformaldehyde (Sigma-Aldrich-P6148), MTT (Sigma-Aldrich-M5655), Trizma (Sigma-Aldrich-T6066), SDS (Sigma-Aldrich-L3771), Uric acid sodium salt (Sigma-Aldrich-U2875), Suberic acid (Sigma-Aldrich-S1885), OPTI-MEM media (Gibco-11058-021), β-Amyloid (Anaspec-AS-60479).

Antibodies-NLRP3 (CST-15101S), ASC (Santa Cruz Biotechnology-SC-22514), CASP1 (Santa Cruz Biotechnology-SC-56036), anti-mIL-1β (R & D biotechnology-AF-401-NA), pCAMKK2 (CST-12818S), PRKAA2 (CST-2532S), pPRKAA2 (CST-2535S), pULK1 (CST-14202S), ULK1 (Santa Cruz Biotechnology-SC-33182), FIP200 (CST-12436S),BECN1 (Santa Cruz Biotechnology-SC-48341), ATG5 (Santa Cruz Biotechnology–SC-133158), ATG7 (Santa Cruz Biotechnology-SC-33211), ATG13 (CST-D4P1K), NF-κB (Santa Cruz Biotechnology-SC-372), pNF-κB (Santa Cruz Biotechnology-SC-136548), MTOR (CST-2972S), pMTOR (CST-5536S),LAMP1 (Santa Cruz Biotechnology-SC-20011), HRP-linked anti-goat IgG (Santa Cruz Biotechnology-SC-2354), siRNA PRKAA2 (CST-6620S), GFAP (CST-80788S), IBA1 (CST-17198S), HRP-linked anti-rabbit IgG (CST-7074S), HRP-linked anti-mouse IgG (CST-7076S), anti-mouse IgG Alexa flour 488 (CST-4408S), anti-mouse IgG Alexa flour 555 (CST-4409S), anti-rabbit IgG Alexa flour 488 (CST-4412S), anti-rabbit IgG Alexa flour 555 (CST-4413S), anti-ACTB (Sigma-Aldrich-A3854), ANTI-LC3B-II (Sigma-Aldrich-L7543), Anti-p62/SQSTM1 (Sigma-Aldrich-P0067).

Kits and reagents-PVDF Membrane (Millipore-ISEQ00010), Precision plus protein markers (Bio-Rad-161-0375), ECL-kit (Millipore-WBKLS0500), Bradford reagent (Bio-Rad-500-0006), FuGENE HD (Promega-E2313), mouse TNF-α ELISA kit (Invitrogen-88-7324-88), mouse IL-6 ELISA kit (Invitrogen-88-7064-88), mouse IL-1β ELISA kit (Invitrogen-88-7013-88), mouse IL-18 ELISA kit (Invitrogen-88-50618-88), Human Aβ42 ELISA kit (Invitrogen-KHB3441)

### Synthesis of fluvoxamine maleate

The synthesis of fluvoxamine maleate was done by a well-established method (Figure 1A) (WO2014178064, US 6,433,225 B1). The scheme for synthesis of fluvoxamine maleate (Scheme 1) and data related to its characterization is given in the supplementary material file.

**Figure 1.**
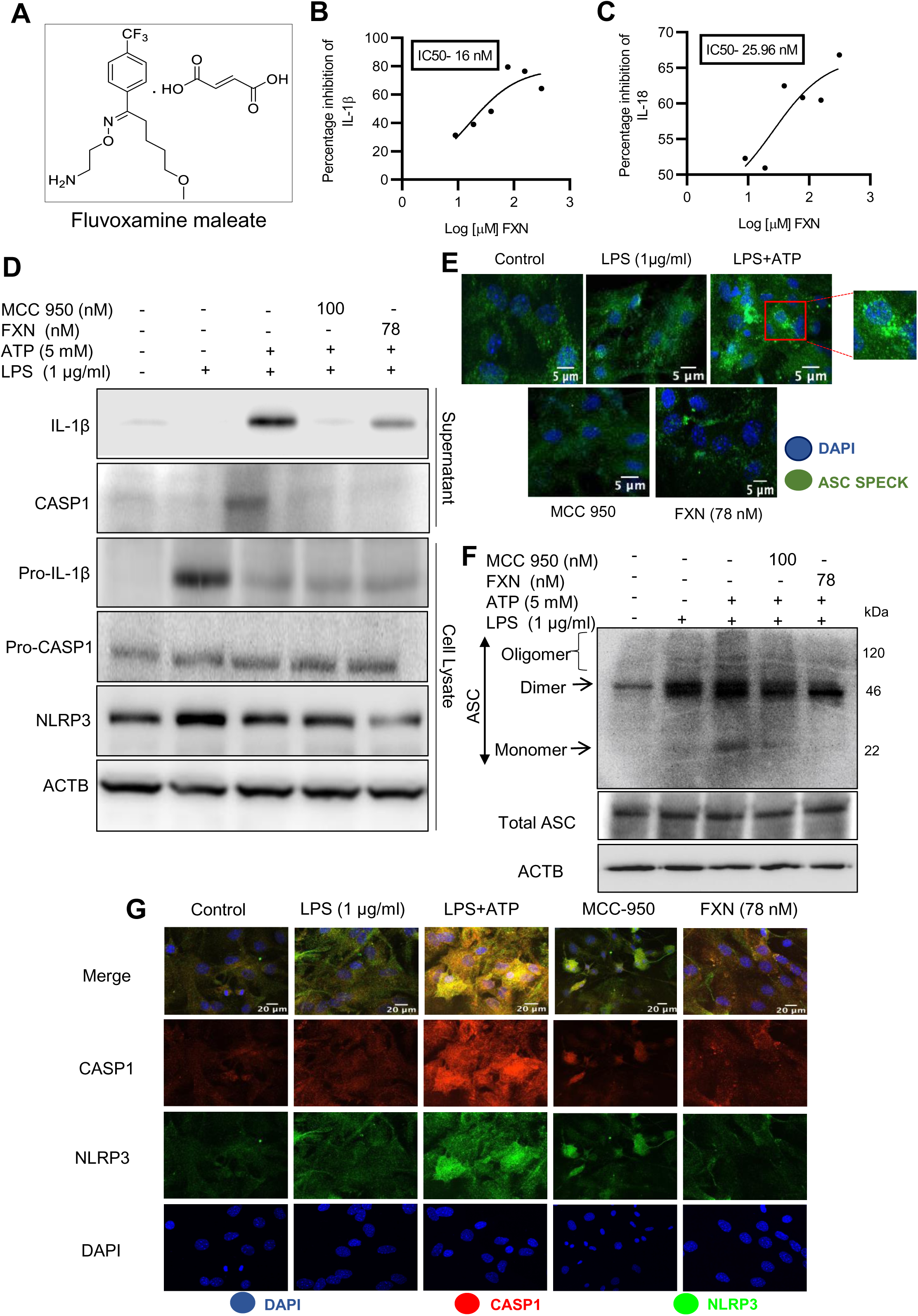
FXN displayed a potent anti-NLRP3 inflammasome activity in primary astrocytes. (A) Structure of Fluvoxamine maleate. (B) and (C) Graphs representing IC_50_ of FXN for IL-1β and IL-18, respectively. The IC50 values were calculated in primary astrocytes after treatment with FXN in the serum free media for 1 h and ATP (5 mM) for 30 minutes. (D) Immunoblots depicting the expression of NLRP3 inflammasome complex proteins including IL-1β, CASP1 and NLRP3. Densitometric analysis of the western blots is provided in Supplementary data (Fig. S1A). (E) Representative confocal images indicating the ASC speck formation in primary astrocytes, which generally form after NLRP3 inflammasome activation, and the graph showing average number of specks is provided in the Supplementary data (Fig. S1B). (F) Western blots showing the ASC oligomerization. Densitometric analysis of immunoblots is given in Supplementary data (Fig. S1C). (G) Representative images showing co-localization of NLRP3 (green) and CASP1 (red) in FXN (78 nM) treated primary astrocytes. The mean fluorescence intensities of NLRP3 and CASP1 is provided in the Supplementary data (Fig S1D). The above-mentioned findings revealed that FXN significantly decreased the release of IL-1β and IL-18 and inhibited the expression of proteins involved in the NLRP3 complex formation including CASP1, NLRP3 and ASC. The data provided are Mean±SD of three independent experiments and the statistical analysis was measured using one-way ANOVA, followed by post-hoc Bonferroni test. The p-value<0.05 was considered to be statistically significant with values assigned as ****p < 0.0001, ***p < 0.001, **p< 0.01, *p< 0.05 and ns= not significant.

## Cell culture

### Primary astrocyte culture

The brain was isolated from four to five days old pups of C57BL/6 mice and cortical hemisphere was treated with trypsin. Following centrifugation, the cells were passed through cell strainer and cultured in low glucose DMEM media and supplemented with 10% FBS. After every 2 days, fresh media was added to the cells. After one week, the shaking was carried out using rocker to separate out the astrocytes from the top layer of the microglial cells. Thereafter, astrocytes were trypsinized and cultured for future experiments under 95% humidity and 5% CO_2_ at 37°C in an incubator chamber.

### Cell line maintenance

The N9 microglial cells were cultured in RPMI-1640 media supplemented with 10% Fetal bovine serum (FBS), penicillin (70 mg/L), streptomycin (100 mg/L) and NaHCO_3_ (3.7 g/L), under constant supply of 5% C0_2_ and 95% humidity at 37°C.

### NLRP3 inflammasome activation in primary astrocytes

In order to activate the NLRP3 inflammasome in primary astrocytes, N9 microglial cells were seeded in 24 well plates and allowed to grow for 24 h. The cells were then treated with LPS (1 μg/ml) and incubated for next 24 h. The LPS activated microglial conditioning media was collected and primary astrocytes were pretreated with the conditioned medium for 24 h. Next day, primary astrocytes were primed with LPS (1 μg/ml) for 4 h and treated with different concentrations of FXN and standard MCC 950 (100 nM) for 1 h in serum free media, followed by ATP (5 mM) stimulation for 30 min.

### Enzyme-linked Immunosorbent Assay (ELISA)

After activation of NLRP3 inflammasome and drug treatments (as mentioned in the previous section), the supernatants were collected and analyzed to assess the levels of IL-1β and IL-18 using ELISA following the manufacturer’s instructions (Invitrogen). For detection of LPS induced pro inflammatory cytokines, TNF-α and IL-6, primary astrocytes were seeded in 24 well plate and pretreated with N9 conditioning media. Then the astrocytes were incubated with LPS for 24 h and the levels of pro-inflammatory cytokines were examined in the supernatants. The levels of cytokines were normalized by dividing the values with total protein content present in the sample. For *in vivo* studies, brain cortex was homogenized in RIPA buffer and centrifuged at 12000 rpm for 20 min at 4°C. The supernatants were collected and analyzed for IL-1β, TNF-α and IL-6 cytokine levels. After completion of 2 months oral dosing, blood was collected from retro-orbital plexus and the Aβ_42_levels in plasma of 5X FAD mice were analyzed using ELISA following the manufacturer’s guidelines (Invitrogen).

### ASC Oligomerization

After drug treatments under NLRP3 inflammasome activation condition, primary astrocytes were lysed in the cold buffer consisting of 1% cocktail, sodium orthovanadate (1 mM), KCL (150 mM), PMSF (0.1 mM), HEPES-KOH (20 mM) and 1% of NP-40. The lysed cells were centrifuged at 330g for 10 min at 4°C and the supernatant was collected to analyze the protein expression by western blotting. The cell pellets were washed with PBS and 500 μL of cold PBS was added to the cells. Suberic acid (2 mM) was added and incubated with the pellets for 30 min at 37°C for cross-linking of ASC proteins. Afterwards the pellets were centrifuged at 330g for 10 min at 4°C and 2X Laemmli buffer was added to the cell pellets. Cell lysates were heated at 95°C for 5 min and proteins were analyzed by western blotting.

### Western blotting

For *in vitro* protein analysis by western blotting, primary astrocytes were seeded in 60 mm petri dishes and treated with FXN and standard MCC 950 under NLRP3 inflammasome activation condition. Cytokines secreted in the media were concentrated using Strataclean resin according to the instructions provided by manufacturer and cells were lysed with RIPA buffer [PMSF (2mM), Na3OV4 (0.5mM), NaF (50mM) and 1% cocktail] and centrifuged at 12,000 rpm for 20 min at 4°C. For *in vivo* studies, hippocampi from 5X FAD mice brain were homogenized and the lysed samples were centrifuged. Supernatants were collected and the protein content was estimated using Bradford’s reagent. For in vitro studies 60 μg protein and for *in vivo* studies 20 μg protein was loaded for separation using SDS PAGE and proteins were transferred to PVDF membrane for 2 h at 4°C. The blots were blocked using 5% BSA or 5% skimmed milk for 1 h at RT and incubated overnight with primary antibody at 4°C. Afterward blots were incubated with HRP-conjugated secondary antibody for 2 h at RT. Chemiluminescent HRP substrate (Millipore) was used for detection of protein bands and visualized by Chemidoc system (Syngene G:BOX chemi XT 4).

### Autophagy flux measurement

Autophagy flux was measured by treating primary astrocytes with different concentrations of FXN in the presence and absence of bafilomycin A1 for 24 h. The protein expression of LC3B-II and SQSTM1 were analyzed by western blotting.

### Confocal microscopy

For confocal microscopic examination of the proteins, primary astrocytes were seeded in 6-well plate over coverslip. After different treatments, cells were washed thrice with PBS and fixed with 4% paraformaldehyde for 15 min. The cells were permeabilized with 0.2% Triton X-100 for 7 min and blocked using blocking buffer (2% BSA and 0.2% Triton X-100) for 1 h. The cells were incubated overnight with primary antibody at 4°C. Next day, the cells were incubated with secondary antibody (Alexa flour 555 or Alexa flour 488) for 1 h at RT and washed thrice with PBS. After counterstaining the nuclei with DAPI, the coverslips were mounted using the mounting media and the images were acquired using Yokogawa CQ1 Benchtop High-Content Analysis System at 40× or 60×.

### A**β** clearance assay

Aβ_42_-HiLyte flour488 peptide was prepared following the manufacturer’s protocol (Anaspec Inc.). Primary astrocytes seeded on coverslips were treated with FXN or rapamycin. After 12 h, 2 μg/ml fluorochrome tagged (Hilyte flour488) Aβ_42_ protein was added to cells for another 12 h. Bafilomycin A1 was given 3 h prior to experiment termination. Cells were washed with PBS and fixed with 4% PFA, afterwards cells were permeabilized with Triton-X 100, and the nuclei were counterstained with DAPI. Slides were prepared and images were taken and analyzed in the Yokogawa CQ1 Benchtop High-Content Analysis System at 40× or 60×.

### Transfection of primary astrocytes with PRKAA2 *siRNA*

Primary astrocytes were grown in 6-well plate for confocal microscopic analysis and 60-mm dishes for western blotting. For transfection of PRKAA2, cells were incubated with OPTI-MEM media and PRKAA2 siRNA in FuGENE HD was added for 24 hours. Transfected primary astrocytes were incubated with microglial conditioning media for 24 h, followed by FXN (78 nM) treatment under NLRP3 inflammasome activation condition and the samples were analyzed by western blotting and confocal microscopy at 60× (Yokogawa CQ1 Benchtop High-Content Analysis System).

### Drug Formulation

For in vivo studies, FXN was formulated in 5% of DMSO, 30% PEG400, 20% PEG200 and 45% distilled water. LPS and ATP were dissolved in PBS.

### Animals and Ethical Clearance

Six months old 5XFAD transgenic mice were used in the study and randomly divided into three groups (n = 5). C57BL6/J mice were used as Control wild type (WT). They were housed under a 12-hour light/dark cycle in a temperature (65–75 °F; ‘18-23 °C) and humidity-controlled (40–60 %) environment, supplied with free access to the food and water (ad libitum). Prior to initiation of study all the animals were acclimatized for one week under standard laboratory conditions, animals were drug naive with no prior procedures performed. All testing were performed from 1 to 4 p.m. Mice were randomized in groups based on their body weights for all behavioral assays and testing were performed by an experimenter blinded to the treatment groups. Total study duration was 2 months. All experiment protocols were approved by the Institutional Animal Ethics Committee (IAEC) (IAEC approval no.-321/82/2/2023), and followed the Committee for Control and Supervision of Experiments on Animals (CCSEA; Ministry of Environment and Forest, Government of India) guidelines for animal care.

### Open-field test

The open field arena box (60 cm x 45 cm x 25 cm) made up of white colored non-reactive plastic was used for all the assessments. The mice were acclimatized to the testing room 30 minutes prior to experiment. Individual mice were placed in the center of the open field arena and allowed to freely explore it for 5 minutes. The arena box was cleaned with a 70% ethanol after every trial. The locomotor activity and exploratory behavior of mice were recorded using a video camera connected to AnyMaze software. Mice were assessed in the open field and the following parameters were recorded automatically: Total distance travelled (cm), average speed (m/s), time spent in the center area (sec), and time spent in the corners (sec).

### Radial arm maze test

The eight-armed radial arm maze (UGO Basile) was used to assess spatial memory behavior. The mice were habituated to the maze for three days before the experiment. Each mouse was trained to reach to the baited arm having a butter cookie as food reward and flag as a visual cue. Training trial was of 3 days. On the memory retention trial day the mouse was placed on the end of one arm (entry arm) and allowed to navigate the baited arm. The movement of mice was tracked with the help of automated AnyMaze software connected with a video tracking camera.

### Neuromuscular coordination test

The neuromuscular coordination of mice was recorded by rotarod. For this assessment mice were trained for three consecutive days with three trials per day. The maximum time allowed on the rotarod during the experiment was 300 seconds. The rotarod was initially set at a speed of 4 rpm at the start of the test and accelerated to 40 rpm over 300 seconds. The latency to fall from the rotarod was recorded for each mouse, and the mean latency to fall for each group was calculated.

### Immunohistochemical analysis of 5XFAD mice brains

After the treatment completion, the hippocampi were isolated from the mice brains and the 10 μm thick tissue-sections were prepared. For IHC, the slides were deparaffinized and histochemical staining was carried out. The sections were incubated with primary antibody (overnight at 4°C under humid conditions after blocking using 2% BSA solution in TBS. For analysis of total Aβ, sections were incubated with anti-Aβ antibody (1-42) D3E10 followed by secondary antibody (Alexa fluor 488) and for localizing reactive astrocytes, anti-GFAP was used followed by secondary antibody (Alexa fluor 555). The alteration in autophagy levels were assessed by examination of LC3B levels in the tissue sections of mice hippocampi. The sections were immuno-stained with LC3B followed by secondary antibody (Alexa fluor 488). The nuclei were counterstained using DAPI. The images were acquired and quantified using Confocal Quantitative Image Cytometer at 40X (Yokogawa CQ1).

### Statistical analysis

Statistical analyses were analyzed using Graph pad prism 9 software. The data shown here are Mean±SD of three independent experiments. The response of independent mice from the group was noted for animal experiments and the mean was calculated. The statistical analysis of data was calculated using one-way ANOVA, followed by post-hoc Bonferroni test. The p-value<0.05 was considered to be statistically significant with values assigned as ****p < 0.0001, ***p < 0.001, **p< 0.01, *p< 0.05 and ns= not significant.

## Results

### FXN displayed a potent anti-NLRP3 inflammasome activity in primary astrocytes

NLRP3 inflammasome being an important player in neuroinflammation, we treated the LPS-primed primary astrocytes with FXN before induction with ATP to analyze the anti-NLRP3 inflammasome activity of FXN. The ELISA results showed that FXN could inhibit NLRP3 inflammasome even at very low concentrations of nine nanomolar, which was evidenced by the measurement of released IL-1β and IL-18 as inflammasome activation products. FXN inhibited the release of IL-1β and IL-18 with IC50 values of 16 nM & 25.96 nM, respectively (Figure 1B and C). Based on the concentration-dependent inhibitory activity of FXN, we chose 78 nM concentration for further detailed studies. We also confirmed the inhibition release of cleaved IL-1β and CASP1 through western blotting of the proteins collected from the supernatant of primary astrocytes treated with FXN (Figure 1D). The analysis of whole cell lysates revealed reduced levels of pro-IL-1β and NLRP3, while pro-CASP1 was unchanged (Figure 1D and Supplementary Figure S1A). Further, the oligomerization of ASC is an important step in the activation of NLRP3 inflammasome. Therefore, we analyzed the effect of FXN on ASC by two methods, including confocal microscopy and western blotting. Both experiments showed similar results with the potent inhibitory effect of FXN on ASC oligomerization (Figure 1E and F and Supplementary Figures S1B and C). We also analyzed the co-localization of CASP1 and NLRP3 in the presence of FXN. Data revealed that FXN and MCC950 (used as a standard) had a similar inhibitory effect on the co-localization of CASP1 and NLRP3 (Figure 1G and Supplementary Figure S1D).

### FXN inhibited the NLRP3 inflammasome through NF-**κ**B pathway

In the initial experiments, we treated the LPS-primed astrocytes with FXN to know whether it could inhibit the assembly of the NLRP3 inflammasome complex. We also wanted to know if it can inhibit the NLRP3 inflammasome at the transcriptional level by inhibiting the NF-κB. Therefore, we treated the cells with FXN before priming them with LPS. We found that FXN could significantly inhibit NLRP3 inflammasome at a concentration of 78 nM contrary to nine nanomolar, observed during FXN treatment post-priming. The ELISA measurement of IL-1β showed an IC50 value of 131.1 nM (Figure 2A). We confirmed the inhibitory effect of FXN on NF-κB by direct and indirect methods. The dwindling levels of NF-κB dependent proinflammatory cytokines TNF-α and IL-6 after treatment with FXN at 78 nM indicated the inhibition of NF-κB in astrocytes (Figure 2B & C). Direct evidence for inhibition of NF-κB pathway came from western blot analysis of NF-κB in the cytosolic and nuclear fraction of astrocytes treated with FXN. The data showed that the movement of NF-κB, p65 from the cytoplasm to the nucleus is hindered in the cells treated with FXN, wherein the levels of NF-κB in the cytoplasm were significantly higher than in the untreated control. On the contrary, its level in the nucleus was significantly reduced (Figure 2D-F).

**Figure 2.**
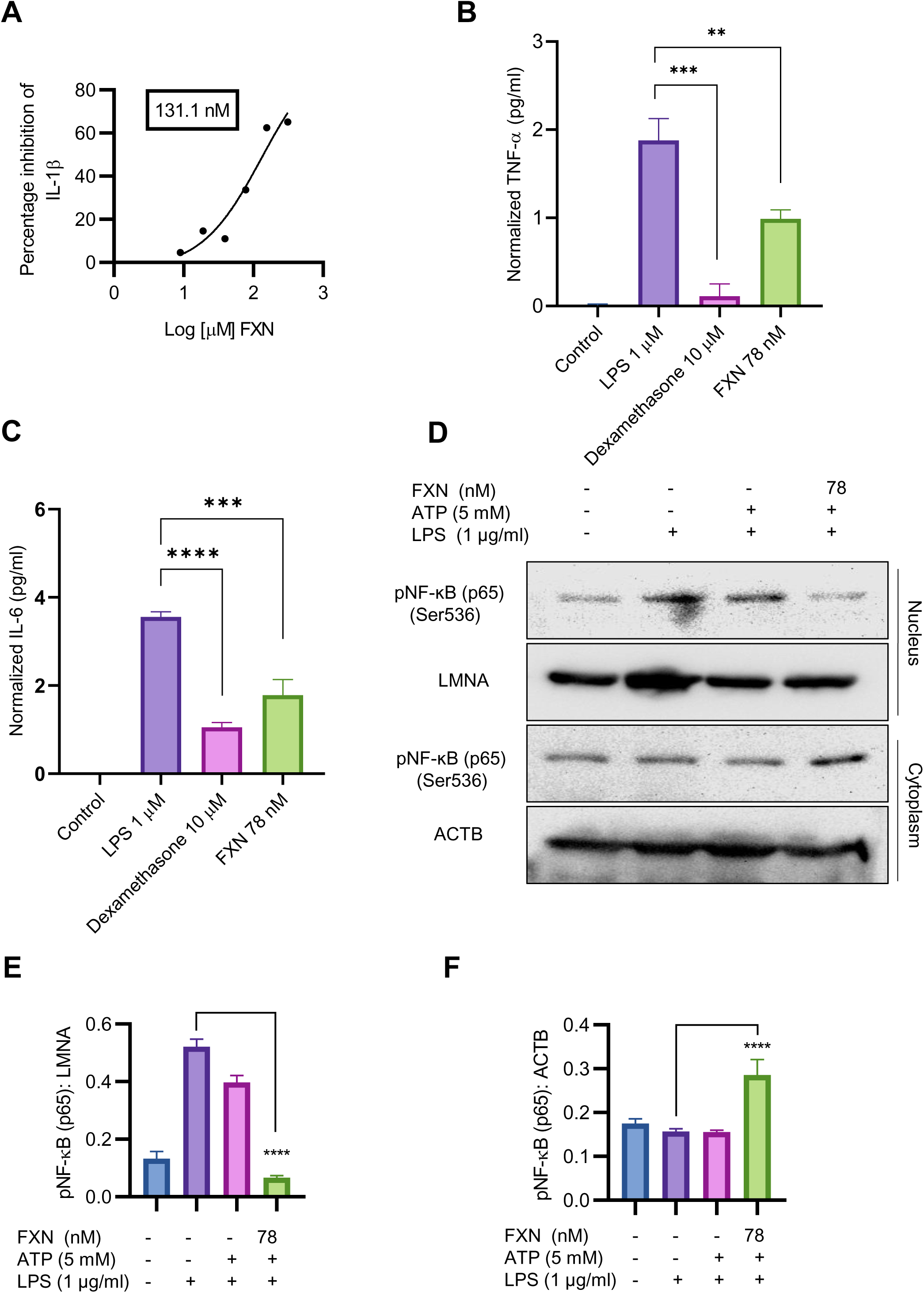
FXN inhibited the NLRP3 inflammasome through NF-κB pathway. The effect of FXN on NF-κB pathway was analyzed by treating the primary astrocytes with FXN, prior to LPS priming and ATP activation of NLRP3 inflammasome. The analysis of pro-inflammatory cytokines regulated by NF-κB pathway was done through ELISA. (A) Graph representing IC50 value of FXN for IL-1β, (B) and (C) Graphs representing the TNF-α and IL-6 levels, respectively, after FXN treatment (78 nM). (D) Analysis of nuclear translocation of NF-κB (p65) through western blotting. LMNA was used as a nuclear marker and ACTB was used as cytosolic marker. (E) and (F) showing densitometric analysis of nuclear and cytosolic fractions of NF-κB (p65), respectively. The data indicates the significant reduction in the NF-κB mediated secretion of TNF-α and IL-6 by FXN treatment via inhibiting the nuclear translocation of NF-κB (p65). The data shown here are Mean±SD of three independent experiments and the statistical analysis was calculated using one-way ANOVA, followed by post-hoc Bonferroni test. The p-value<0.05 was considered to be statistically significant with values assigned as ****p < 0.0001, ***p < 0.001, **p< 0.01, *p< 0.05 and ns= not significant.

### FXN induced the autophagy under the inflammatory conditions in primary astrocytes

After confirming the inhibitory effect of FXN on NLRP3 inflammasome, we wanted to know its inhibition mechanism. Therefore, we focused on autophagy as a major mechanism regulating inflammasome activity. We found that FXN could induce autophagy at a low concentration of 39 nM; however, its effect was enhanced with the increasing concentration (Figure 3A and Supplementary Figure S2A). Further, to confirm the completion of autophagy, we calculated the autophagy flux by using LC3B-II and SQSTM1 expression after treatment of astrocytes with FXN (78 nM) in the presence and absence of end-stage autophagy inhibitory bafilomycin A1. We found an autophagy flux at all the tested time points through 24 h of astrocyte treatment with FXN (Figure 3B and Supplementary Figure S2B). Induction of autophagy by FXN was also confirmed by confocal microscopy analysis of co-localization of LC3B-II and LAMP1 in primary astrocytes. The cells treated with FXN and rapamycin showed a significant amount of co-localization of these proteins, while in the presence of bafilomycin A1, FXN-treated cells did not show any observable co-localization of LC3B-II and LAMP1 (Figure 3C and D). After confirming the induction of autophagy by FXN, we checked if it can induce autophagy in primary astrocytes under inflammatory conditions where the cells have been challenged with LPS and ATP. We found that the treatment of astrocytes with FXN before ATP induced the expression of LC3B-II, while SQSTM1 was downregulated, thus confirming the induction of autophagy (Figure 3E and Supplementary Figure S2C). We further analyzed the expression of various proteins involved in the induction of autophagy. We found that FXN induced autophagy through CAMKK2 mediated induction of PRKAA2 pathway leading to activation of various proteins including pPRKAA2 (Thr 172), pULK (Ser 317), ATG13, FIP 200, BECN1, ATG5 and ATG7 and downregulation of pMTOR (Figure 3F and supplementary Figures S2C).

**Figure 3.**
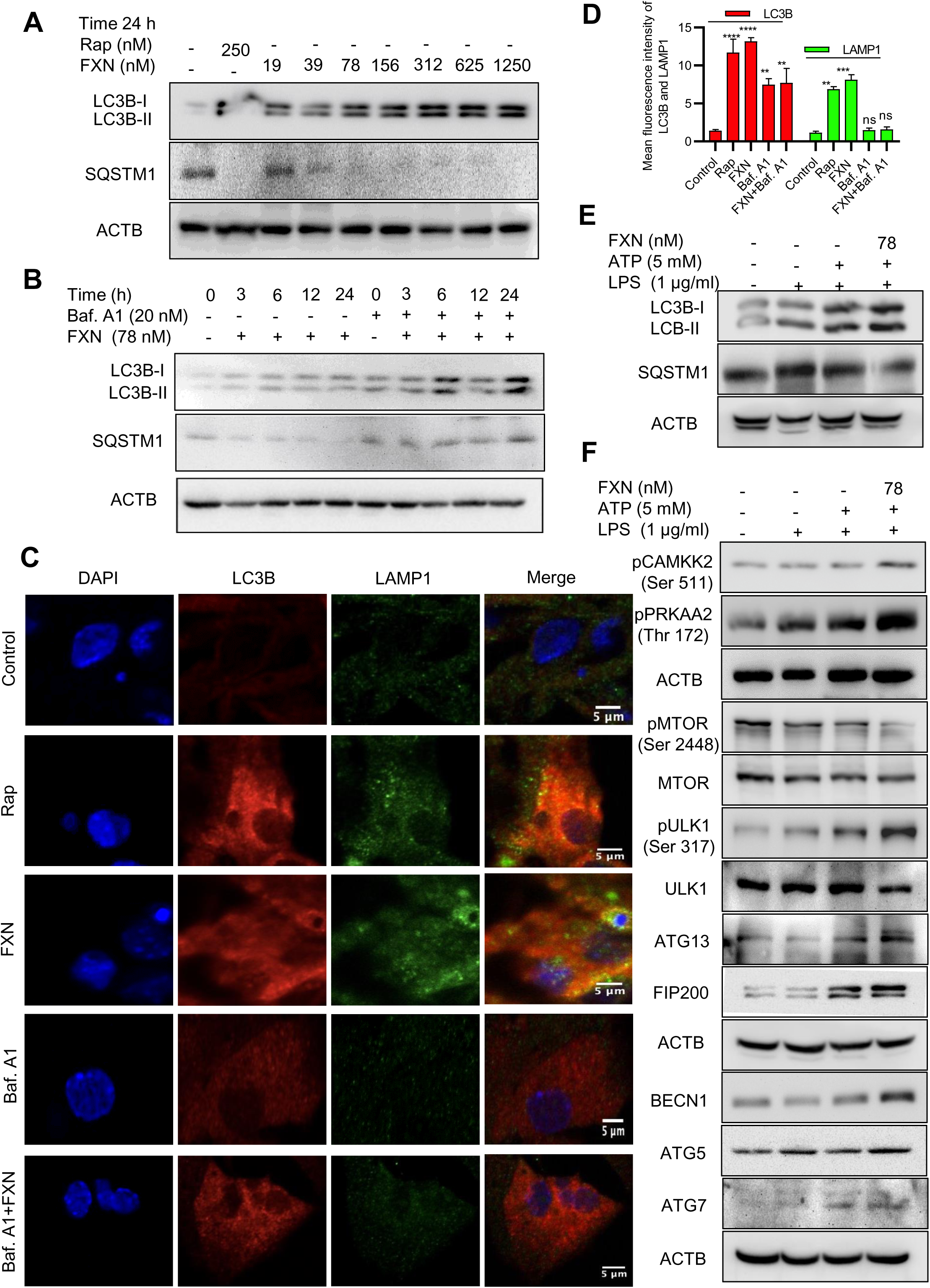
FXN induced the autophagy under the inflammatory conditions in primary astrocytes. (A) Immunoblots showing concentration dependent effect of FXN and rapamycin on the expression of LC3B-II and SQSTM1 after 24 h treatment in primary astrocytes. Densitometric analysis is given in Supplementary data (Fig. S2A). (B) Immunoblots showing the time dependent effect of FXN (78 nM) on autophagic flux in the presence and absence of bafilomycin A1 in primary astrocytes. Densitometric analysis is provided in Supplementary (Fig. S2B). (C) Representative images showing the effect of FXN (78 nM) on co-localization of LAMP1 (green) and LC3B (red) in primary astrocytes. The increased colocalization of LAMP1 and LC3B after FXN treatment indicates the induction of autophagy. (D) Graph representing the mean fluorescence intensities of LAMP1 and LC3B. (E) Immunoblots showing the effect of FXN (78 nM) on LC3B-II and SQSTM1 under NLRP3 inflammasome activation conditions in primary astrocytes. Densitometry of immunoblots is provided in Supplementary Data (Fig. S2C). (F) Immunoblots showing the effect of FXN on pMTOR (Ser 2448) and other autophagic proteins involved in the initiation, elongation, maturation and fusion of autophagosome [pCAMKK2 (Ser 511), pAMPK (Thr 172), pULK1 (Ser 317), ATG 13, FIP 200, BECN1, ATG5 and ATG7] in primary astrocytes. The data provided are Mean±SD of three independent experiments and the statistical analysis was performed using one-way ANOVA, followed by post-hoc Bonferroni test. The p-value<0.05 was considered to be statistically significant with values assigned as ****p < 0.0001, ***p < 0.001, **p< 0.01, *p< 0.05 and ns= not significant.

### FXN inhibited NLRP3 inflammasome by inducing autophagy

For the confirmation of the involvement of autophagy in FXN-mediated inhibition of NLRP3 inflammasome, we inhibited the autophagy through genetic and pharmacological methods. We pretreated the primary astrocytes with *siPRKAA2* and analyzed the expression of autophagic and NLRP3 inflammasome-related proteins. The inhibition of PRKAA2 led to the reversal of the autophagic effect of FXN, which was reflected in the form of a marked decrease in the expression of LC3B-II and BECN1, while SQSTM1 expression was significantly increased (Figure 4A and Supplementary Figure S3A). Similarly, the inhibition of NLRP3 inflammasome by FXN was also reversed in the cells pretreated with *siPRKAA2*. The cleaved IL-1β levels were significantly increased, whereas the reduction in expression of NLRP3 was also reversed in the cells treated with FXN in the presence of *siPRKAA2* (Figure 4A and Supplementary Figure S3A). We also confirmed these findings by using confocal microscopy, where the co-localization of CASP1 and NLRP3 was restored to the level present in cells primed with LPS and ATP when they were treated with FXN along with knockdown of PRKAA2 by using siRNA (Figure 4B and Supplementary Figure S3B). The knockdown of PRKAA2 was confirmed by using western blotting, where the expression of PRKAA2 and pPRKAA2 was found to be significantly reduced (Figure 4C). We found that a similar kind of reversal of the inhibitory activity of FXN against NLRP3 inflammasome could be achieved if the autophagy is inhibited by using the end-stage pharmacological inhibitor bafilomycin A1. The cleavage of IL-1β was almost completely inhibited, while the expression of NLRP3, NF-κB (p65) along with autophagy markers LC3B-II, SQSTM1 was restored when the cells were treated with FXN in the presence of bafilomycin A1 (Figure 4D and Supplementary Figure S3C)

**Figure 4.**
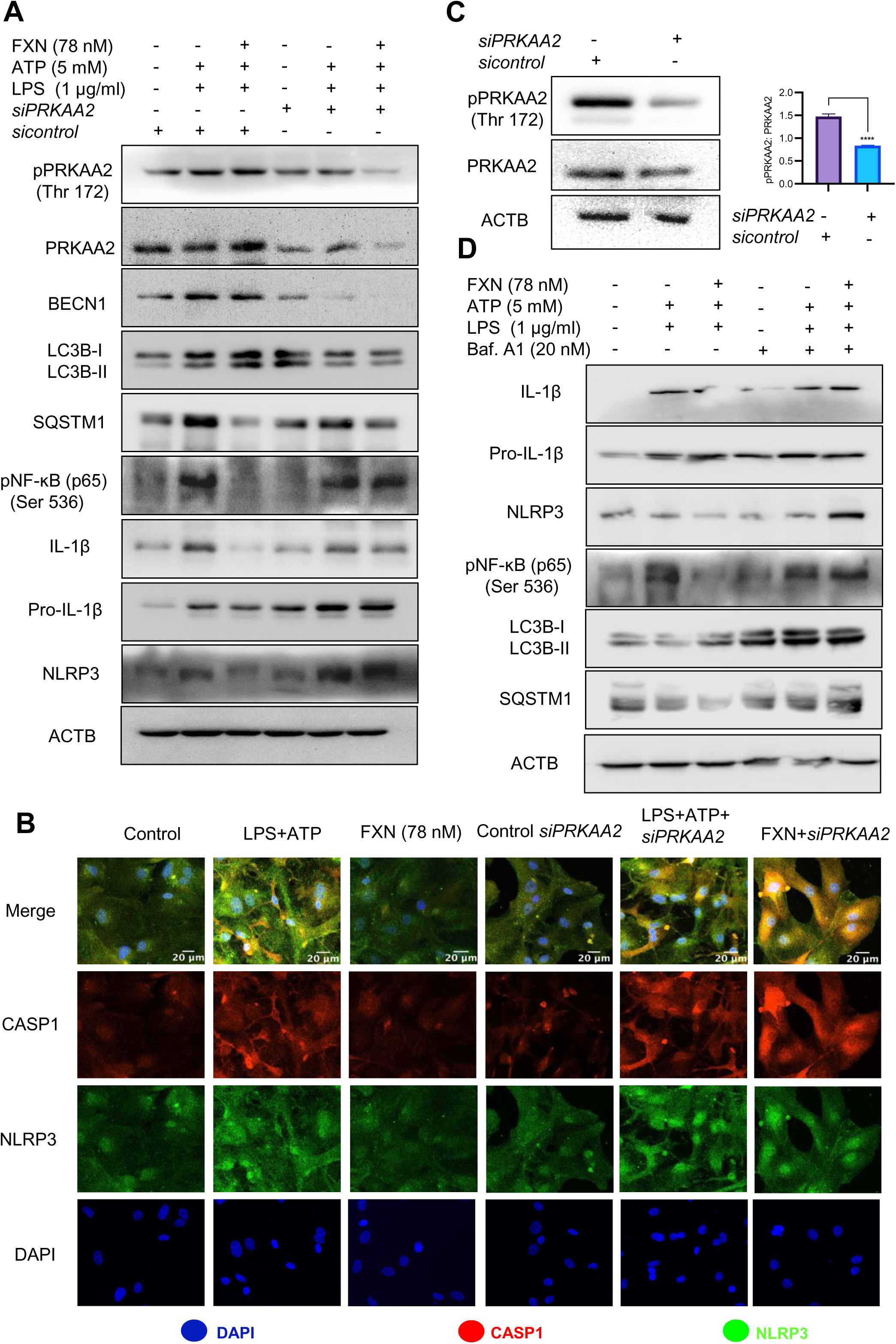
FXN inhibited NLRP3 inflammasome by inducing autophagy. To investigate the involvement of autophagy in the FXN mediated inhibition of NLRP3 inflammasome, autophagy was genetically and pharmacologically inhibited by *siPRKAA2* and bafilomycin. A1 respectively in primary astrocytes under inflammasome activation conditions. (A) Western blots indicating the effect of PRKAA2 knockdown on the expression levels of pPRKAA2 (Thr 172), BECN1, LC3B-II, SQSTM1, NF-κB (p65), IL-1β and NLRP3. Densitometric analysis is given in Supplementary data (Fig. S3A). (B) Representative images indicating the colocalization of NLRP3 and CASP1 under aforementioned conditions (genetic knockdown of PRKAA2 by *siPRKAA2*) after FXN treatment. The mean fluorescence intensities of NLRP3 and CASP1 are provided in the Supplementary data (Fig. S3B). (C) Immunoblots depicting the PRKAA2 expression in primary astrocytes following treatment with the *siPRKAA2,* in comparison to mock *siRNA* and their densitometric analysis. (D) Immunoblots showing the effect of FXN on expression levels of IL-1β, NLRP3, NF-κB (p65), LC3B-II and SQSTM1 after pharmacological inhibition of autophagy using bafilomycin A1. Densitometric analysis is given in Supplementary data (Fig. S3C). The data represent Mean±SD of three independent experiments and the statistical analysis of data was analyzed using one-way ANOVA, followed by post-hoc Bonferroni test. The p-value<0.05 was considered to be statistically significant with values assigned as ****p < 0.0001, ***p < 0.001, **p< 0.01, *p< 0.05 and ns= not significant.

### FXN cleared A**β**_42_ in primary astrocytes by inducing autophagy

To confirm whether the autophagy induced by FXN can help in the clearance of Aβ_42_, we treated the cells for a total of 24 h with FXN, including 12 h treatment with hilyte fluor 488 tagged Aβ_42_. The confocal analysis revealed that the cells treated with FXN and rapamycin were almost completely devoid of green fluorescence indicating clearance of Aβ_42_ in comparison to control cells. Further, 3 h treatment of cells with autophagy inhibitor bafilomycin A1 prior to termination of the experiment completely repressed the clearance of Aβ_42_, as indicated by the same level of green fluorescence as that of control cells (Figure 5A and C). The involvement of autophagy in the clearance of Aβ_42_ by FXN was further confirmed when we inhibited the autophagy by using *siRNA* against PRKAA2. The clearance of Aβ_42_ by FXN in the presence of *siPRKAA2* was similarly stopped as we observed with bafilomycin. A1 (Figure 5B and D).

**Figure 5.**
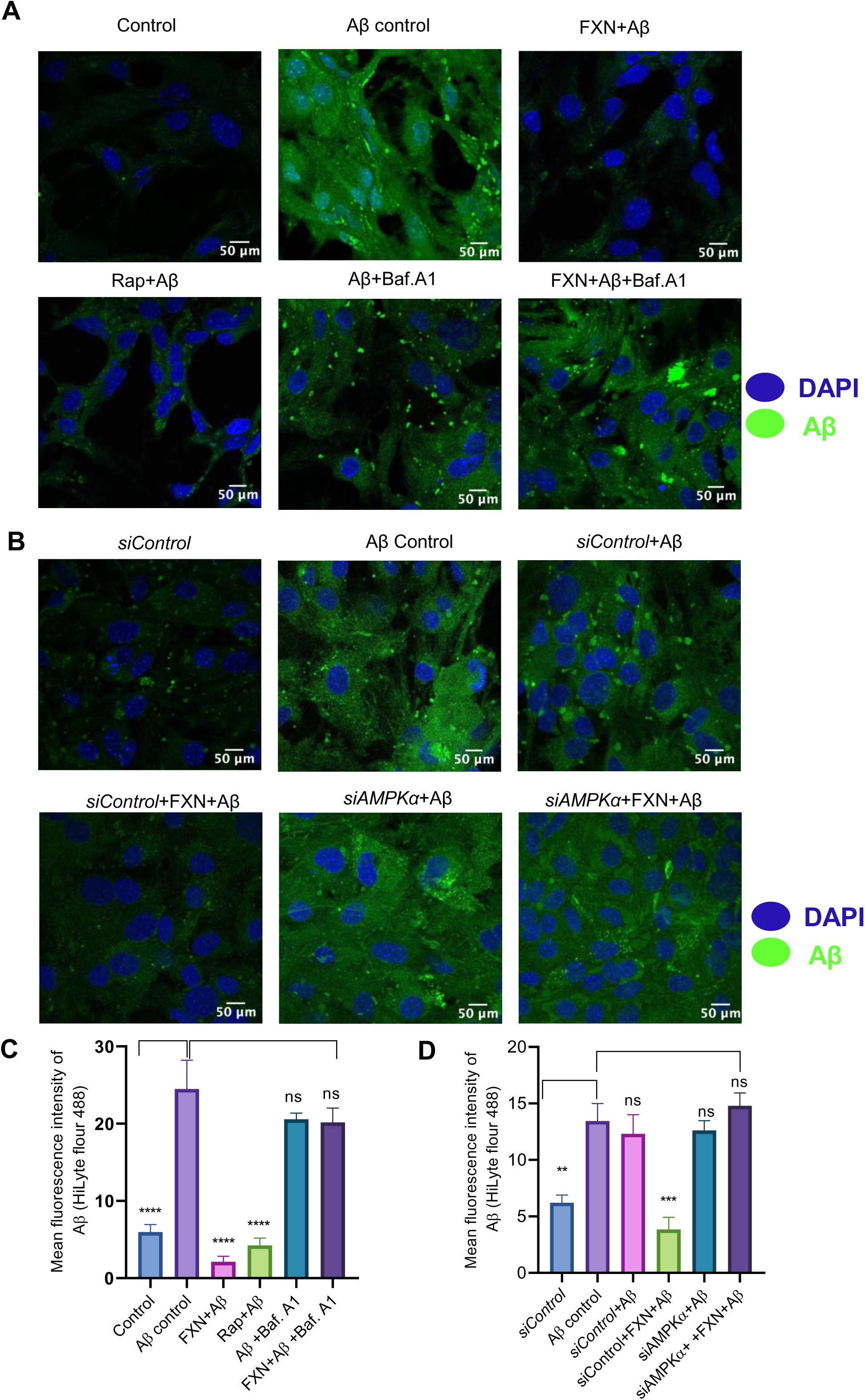
FXN cleared Aβ_42_ in primary astrocytes by inducing autophagy. (A) Representative images showing the effect of FXN, in the presence Baf. A1, on the intracellular deposition of Aβ in primary astrocytes treated with fluorescently tagged (HiLyte 488) Aβ peptide. Rapamycin was used as a standard autophagy inducer. (B) Representative images showing the effect of FXN on intracellular deposition of Aβ in primary astrocytes transfected with *siPRKAA2*. (C) Graph representing the mean fluorescence intensity of images shown in Fig. 5A. (D) Graphical representation of mean fluorescence intensity of images shown in Fig. 5B. The decrease in the fluorescence intensity of Aβ in FXN treated cells shows its potential to clear deposited Aβ and the reversal in the fluorescence intensities of Aβ where the primary astrocytes were treated with Baf. A1 or transfected with *siPRKAA2,* indicates the involvement of autophagy in FXN mediated Aβ clearance. The data shown here are Mean±SD of three independent experiments and the statistical analysis of data was calculated using one-way ANOVA, followed by post-hoc Bonferroni test. The p-value<0.05 was considered to be statistically significant with values assigned as ****p < 0.0001, ***p < 0.001, **p< 0.01, *p< 0.05 and ns= not significant.

### FXN improved working memory, exploratory behavior and neuromuscular coordination in 5XFAD mice

After confirming the mechanism of inhibition of NLRP3 inflammasome by FXN, we wanted to know if its dual pharmacological activity can improve Alzheimer disease pathology. Therefore, we treated the 6-month-old 5XFAD mice for two months and analyzed multiple behavior parameters. In the radial arm maze test, the untreated 5XFAD mice displayed significant deterioration of neuronal functions in comparison to wild-type control (Figure 6A). However, the groups treated with FXN (5 and 10 mg/kg) took significantly less time and traveled a reduced distance in finding the baited arm and spent more time in the baited arm apart from fewer entries into non-baited arms in comparison to untreated 5XFAD mice, indicating improvement in the working memory (Figure 6B-E). In the open field test, untreated 5XFAD mice spent significantly more time in the center than in corners and showed little interest in exploring the surroundings (Figure 6F). However, the mice treated with FXN covered a significantly higher distance at greater speed and showed normal behavior by spending more time in the corners than in the center (Figure 6G-K). We also observed a significant improvement in the neuromuscular coordination in the mice treated with FXN, which showed delayed latency to fall than the untreated 5XFAD mice (Figure 6L). In all the behavior parameters studied, the mice treated with FXN behaved more like wild-type animals than the untreated 5XFAD control group.

**Figure 6.**
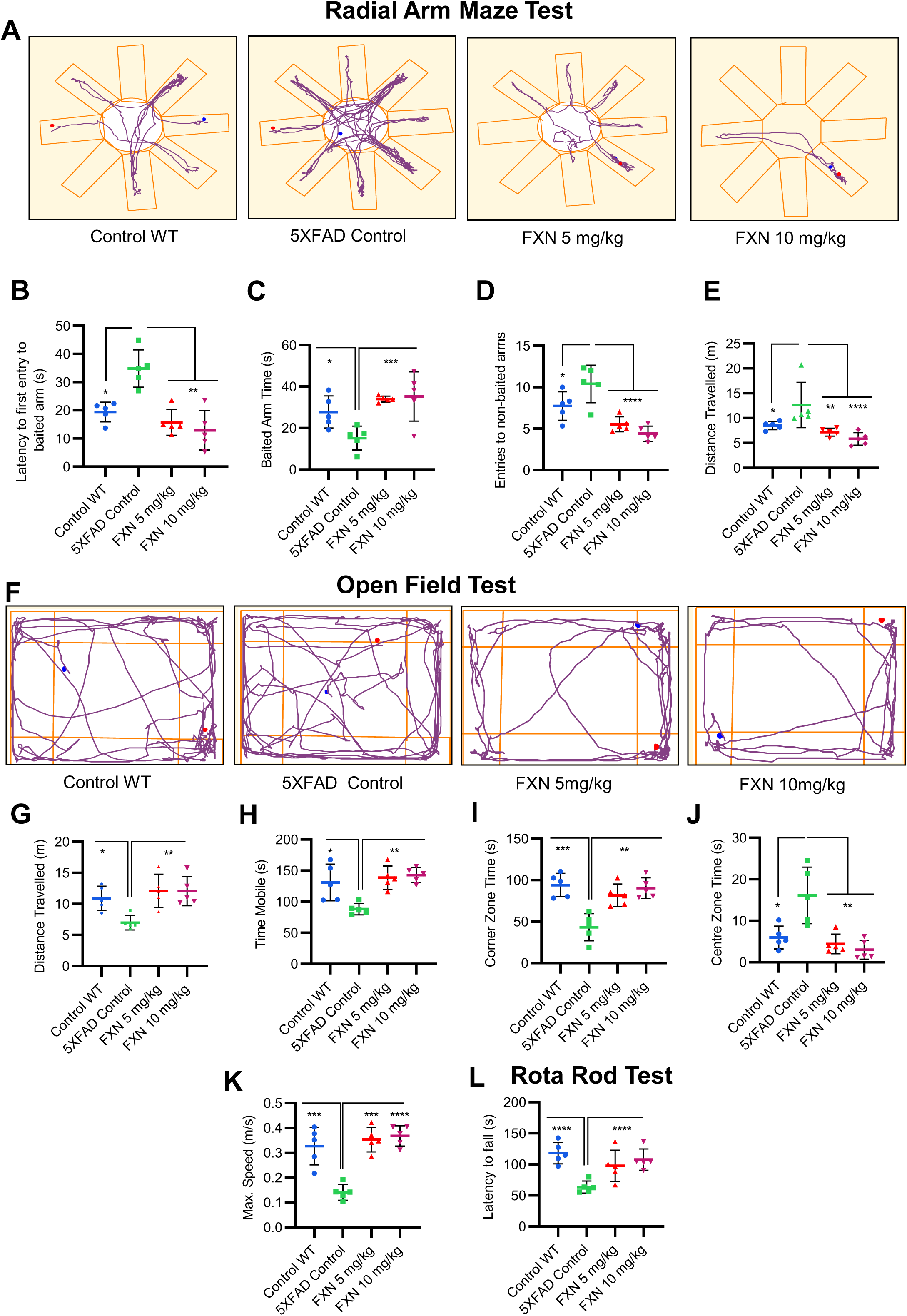
FXN improved working memory, exploratory behavior and neuromuscular coordination in 5XFAD mice. The effect of FXN (5 and 10 mg/kg) on cognitive behavior was assessed post 2 months dosing in Alzheimer (5XFAD) mice model (n=5 per group). Vehicle 5XFAD control mice were compared with wild-type C57BL/6J mice of same age. Radial Arm Maze Test: (A) Representative images showing track plots of radial arm maze post five days training. The different parameters were assessed such as (B) Latency to first entry to baited arm, (C) Time spent in baited arm, (D) Entries to non-baited arms and (E) Distance travelled by mice. (F) Images depicting the track plots obtained from open field test. Various parameters were investigated such as (G) Distance travelled by mice, (H) Mobility time, (I) Time spent in the Centre zone, (J) Corner zone time and (K) Maximum speed. The latency to fall time of mice from rotarod was assessed by using rotarod test. (L) Graph representing the latency to fall time. Automated camera was used to capture the movement of mice in radial arm maze and open field test and all the activities were measured by using AnyMaze software. The statistical analysis of data was measured using one-way ANOVA, followed by post-hoc Bonferroni test. The p-value <0.05 was considered to be statistically significant with values assigned as ****p < 0.0001, ***p < 0.001, **p< 0.01, *p< 0.05 and ns= not significant.

### FXN reduced the A**β**_42_ in the hippocampus and plasma of 5XFAD mice

Improvement in neuronal health of 5XFAD mice after treatment with FXN led us to investigate the underlying cause for such change. Therefore, through immunohistochemistry (IHC), we checked the levels of Aβ_42_ in the hippocampus after two months of treatment with FXN. We found a significant decline in the levels of Aβ_42_ in the hippocampus of both groups of mice treated at 5 and 10 mg/kg, respectively compared to the control group, as indicated by reduced green fluorescence (Figure 7A). The average number of Aβ_42_ plaques, skeletal length of Aβ_42_ plaques, branch count and circumference of Aβ_42_plaques were significantly reduced in FXN treated groups as compared to 5XFAD control group (Figure 7B-E). Aβ_42_ is strongly related to activation of astrocytes, we therefore checked if reduced levels of Aβ_42_ had any impact on the reactive phenotype of astrocytes. The analysis of GFAP in the vicinity of Aβ_42_ revealed that as the levels of Aβ_42_were depleted after treatment with FXN, the expression of GFAP reduced proportionately, as indicated by reduced red color fluorescence (Figure 7A and F). Further, the reduction of Aβ_42_levels in the brain also led to reduced circulatory levels of Aβ_42_ in the blood plasma of 5XFAD mice treated with FXN in comparison to untreated control mice, as observed through ELISA (Figure 7G).

**Figure 7.**
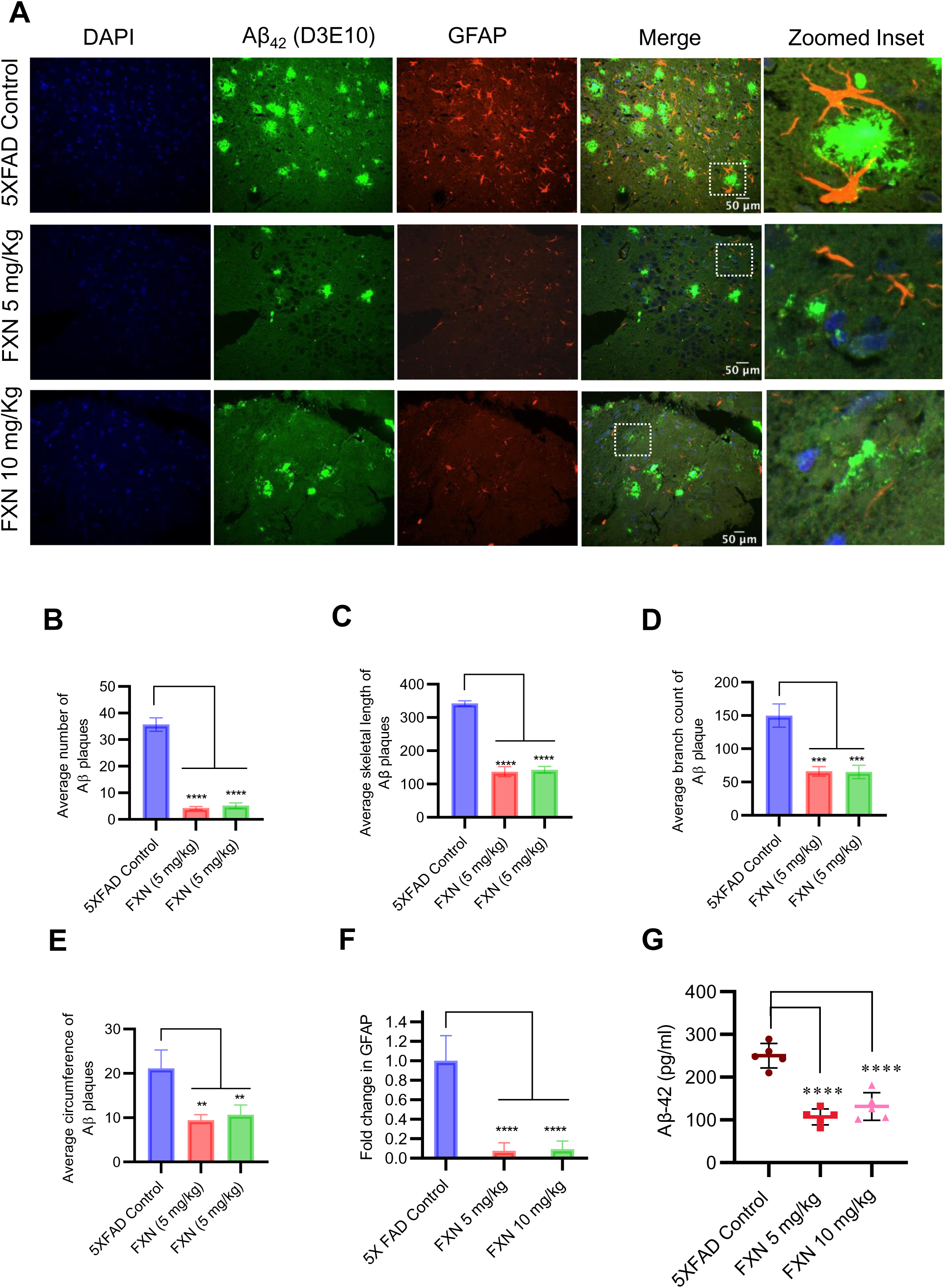
FXN reduced the Aβ_42_ in the hippocampus and plasma of 5XFAD mice. Immunostaining post FXN treatment (5 and 10 mg/kg for 2 months), the formalin-fixed paraffin embedded (FFPE) tissues of brain hippocampi were immuno-stained with D3E10 for detection of total Aβ (green), GFAP for activated astrocytes (red) and the nuclei were counterstained with DAPI (blue). (A) Representative images showing co-localized Aβ plaques and reactive astrocytes in hippocampi of FXN treated mice (n=5 per group). Graphs showing (B) Average number of Aβ plaques (C) Average skeletal length of Aβ plaques (D) Average branch count of Aβ plaques (E) Average circumference of Aβ plaques (G) The plasma levels of Aβ-42 in mice measured using ELISA. The Aβ load and astrogliosis in FXN treated 5XFAD mice was found to significantly reduced. Confocal microscopic images were examined using cell pathfinder software. The statistical analysis was performed using one-way ANOVA, followed by post-hoc Bonferroni test. The p-value <0.05 was considered to be statistically significant with values assigned as ****p < 0.0001, ***p < 0.001, **p< 0.01, *p< 0.05 and ns= not significant.

### FXN ameliorated the AD pathology in 5XFAD mice through autophagy-mediated reduction in amyloid beta levels and neuroinflammation

Based on the effect of FXN on in vitro and in vivo clearance of Aβ_42_, we hypothesized that autophagy induced by FXN is responsible for the reduced level of Aβ_42_ in the hippocampi of 5XFAD mice brains. Therefore, to test the hypothesis, we analyzed the expression of LC3B-II in the hippocampi and found a highly significant increase in its levels in the mice treated with FXN (Figure 8A). Further, analysis of expression of various autophagic proteins including pCAMKK2 (Ser 511), pPRKAA2 (Thr 172), pMTOR (Ser 2448), LC3-IIB, pULK1 (Ser 317), BECN1, and ATG5 revealed a significant upregulation of these proteins in the hippocampi of mice treated with FXN in comparison to untreated mice, while the effect of FXN at 10 mg/kg was more pronounced as compared to the dose of 5 mg/kg (Figure 8B and Supplementary Figures S4A).

**Figure 8.**
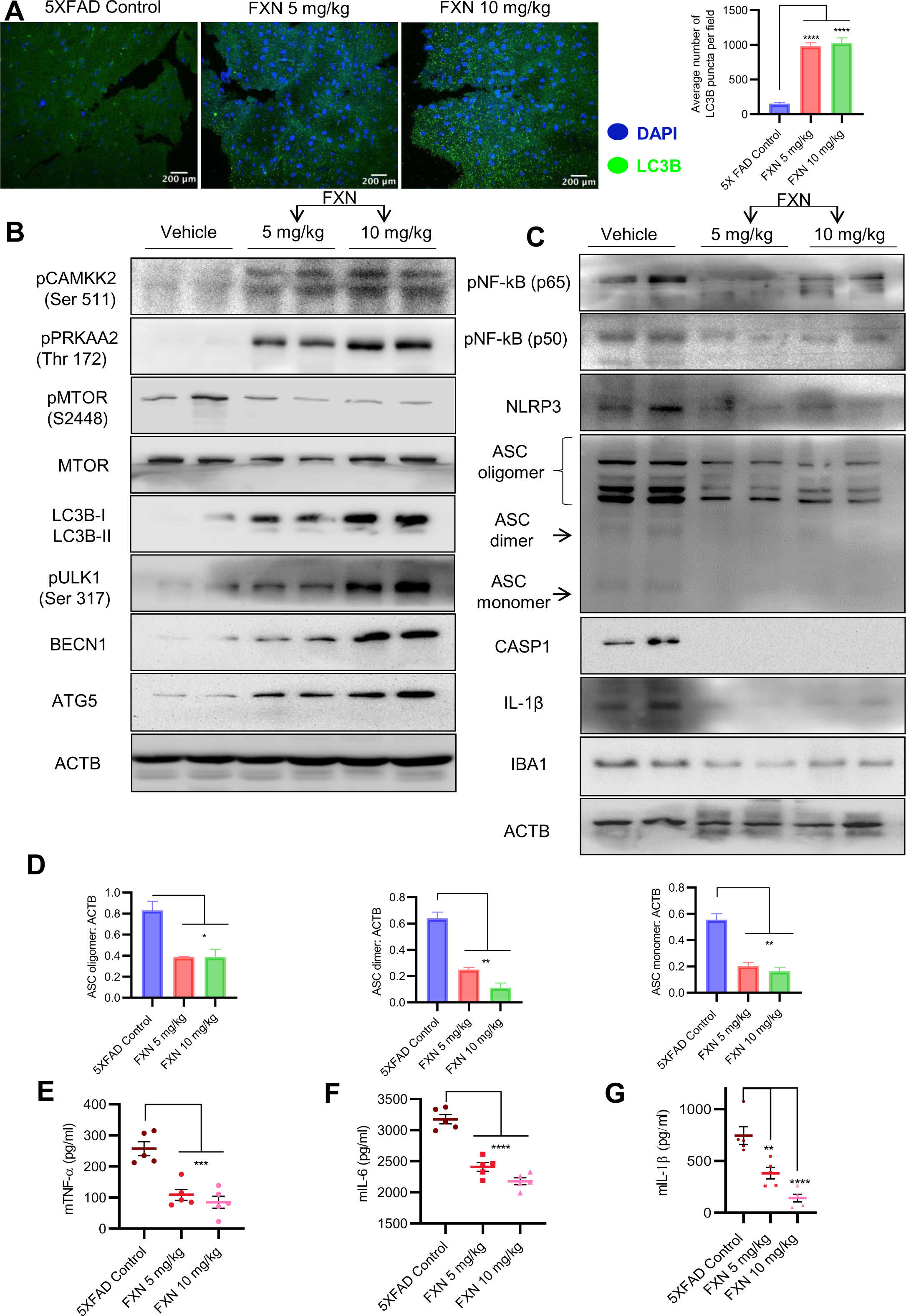
FXN ameliorated the AD pathology in 5XFAD mice through autophagy-mediated reduction in amyloid beta levels and neuroinflammation. (A) Representative confocal images showing LC3B (green) in hippocampi sections of FXN treated mice (5 and 10 mg/kg) (n=5 per group) and the graph representing the average number of LC3B puncta from total of 20 images obtained from each group. (B) Immunoblots showing CAMKK2 mediated autophagy induction via PRKAA2 pathway, indicated by significant increase in the expression of pCAMKK2 (Ser 511), pPRKAA2 (Thr 172), LC3B-II, pULK (Ser 317), BECN1 and ATG5, and a decreased expression of pMTOR (Ser 2448) in hippocampi of FXN treated mice. Densitometric analysis of the western blots is provided in Supplementary data (Fig. S4A). (C) Immunoblots depicting the decreased expression of neuroinflammatory markers NF-κB (p50), NF-κB (p65), NLRP3, ASC oligomer, ASC dimer and ASC monomer, CASP1, IL-1β and IBA1 in hippocampi of FXN treated mice. (D) Densitometric analysis showing decreased levels of ASC oligomer, dimer and monomer forms in FXN treated mice. The densitometry of other western blots is provided in Supplementary data (Fig. S4B). Graphs depicting the levels of pro-inflammatory cytokines in brain cortex of FXN treated 5XFAD mice (E) TNF-α, (F) IL-6 and (G) IL-1β. The samples were compared by using one-way ANOVA, followed by post-hoc Bonferroni test. The p-value <0.05 was considered to be statistically significant with values assigned as ****p < 0.0001, ***p < 0.001, **p< 0.01, *p< 0.05 and ns= not significant.

The analysis of pro-inflammatory proteins in the hippocampi also supported the anti-inflammatory effect of FXN. We found that treatment of mice with FXN significantly reduced the levels of both NF-κB, p50 and p65 subunits confirming its effect on the most important inflammatory pathway. Further analysis of NLRP3 inflammasome proteins showed significant down-regulation of NLRP3, which significantly reduced ASC oligomerization and cleaved CASP1 and IL-1β in the mice treated with FXN (Figure 8C and D and Supplementary Figures S4B). The reduction of inflammation in the hippocampi was indicated by reduced levels of microglial marker protein IBA1 (Figure 8C and Supplementary Figures S4B). We also observed a significant decline in the levels of pro-inflammatory cytokines TNF-α, IL-6 and IL-1β, which were measured in the cortex region of the brain by using ELISA (Figure 8E-G). These data emphasized a potent effect of FXN on the pathology of AD in 5XFAD mice.

## Discussion

Alzheimer disease progresses with a complex pathology. However, most cases of AD are accompanied by the presence of Aβ_42_ plaques and neuroinflammation where NLRP3 inflammasome plays a significant role [2, 21–23]. There are multiple types of damage associated molecular patterns (DAMPs) that can upregulate NLRP3 inflammasome in the brain, including Aβ, TAU NFTs, and ATP [24–27]. All these DAMPs contribute to chronic inflammation and AD pathology. We initiated this study focusing on finding NLRP3 inflammasome inhibitors that can be used to mitigate AD pathology. Our preliminary studies found that fluvoxamine maleate (FXN) is a potent inhibitor of NLRP3 inflammasome with an inhibitory concentration in the nanomolar range in ATP-induced primary astrocytes. US FDA had approved FXN for the treatment of obsessive-compulsive disorder (OCD) in 1994. Its proposed mechanism against OCD is through the inhibition of serotonin reuptake transporter (5-HT) [28–30]. However, our studies revealed new targets of FXN that can be effectively exploited against AD. Additionally, being an approved drug for CNS disease, FXN gave us an additional advantage to be used for AD studies without being worried about its safety and crossing of the blood-brain barrier (BBB). For detailed studies on the effect of FXN on neuroinflammation, we continued working on primary astrocytes because of their emerging role in AD and other neurodegenerative diseases. We found that FXN downregulated the expression of NLRP3 and pro-IL1β along with reduced oligomerization of ASC leading to poor assembly of NLRP3 inflammasome and thus cleavage of CASP1. These data indicated that FXN works on both the steps of NLRP3 inflammasome activation to inhibit its assembly. This assumption was confirmed when we observed a significant reduction in the nuclear translocation of NF-κB, p65 and inhibition of pro-inflammatory cytokines TNF-α and IL-6, which depend on it. These data emphasized that FXN has a wider anti-inflammatory spectrum that targets multiple inflammation-related pathways. These findings are important as inflammation is the key driver in neuronal cell death during AD [31, 32].

After confirming the anti-inflammatory activity of FXN, we intended to know how the upstream mechanisms of two important molecular pathways were inhibited by FXN. Therefore, we explored the possible involvement of autophagy in the FXN mediated downregulation of inflammation. Autophagy can regulate inflammation through multiple ways including recycling of proteins, and organelles like mitochondria, clearance of protein aggregates and other DAMPs, invading microorganisms and PAMPs associated with them, etc. [33–35]. We found that FXN is a very potent inducer of autophagy, which works in the same concentration range, which was effective against inflammation. We further observed that it can also induce autophagy under the inflammatory conditions generated by using LPS and ATP. The treatment of astrocyte with FXN under inflammatory conditions showed a significant decline in the expression of NF-κB and other proteins associated with activation of NLRP3 inflammasome. This was an interesting observation. However, we wanted to be sure about the involvement of autophagy in the anti-inflammatory effects of FXN. Therefore, we inhibited the autophagy by both pharmacological and genetic means and found that all the anti-inflammatory effects of FXN were reversed, which was evidenced by re-instatement of expression level of inflammatory proteins to the pre FXN treatment levels.

Apart from being an important regulator of inflammation, the key physiological role of autophagy is to remove protein aggregates [35]. The failure of autophagy to clear proteins aggregates is strongly linked to number of pathological conditions including AD, where it helps to maintain the physiological levels of Aβ along with other clearance mechanisms [36]. We found that FXN could also help in the clearance of Aβ by inducing autophagy in primary astrocytes. This was further confirmed when the clearance of Aβ was completely reversed in the presence of *siPRKAA2*.

These findings encouraged us to validate the efficacy of FXN to ameliorate the AD pathology in transgenic 5XFAD mice. Therefore, we treated the 5XFAD mice for two months with FXN at two different doses. At the end of treatment, the analysis of different behavior parameters revealed that the mice treated with both doses of FXN had significantly improved brain health. The data from different parameters studied during the radial arm maze test showed a marked improvement in the working memory of mice. The hippocampus is the most important part of the brain that processes working memory and is severely affected during AD, as was observed in untreated control mice. Further, the exploratory behavior of FXN-treated mice was also significantly improved and it appeared more like non-demented animals. They spent more time in corners, traveled more distances at higher speeds, etc. The neuromuscular coordination was also found to be similarly improved in mice treated with FXN. All these parameters clearly indicated the improvement of brain health after treatment with FXN. These data made us to explore molecular mechanisms behind this improvement in brain health, particularly memory behavior. Therefore, we isolated the hippocampi of these mice and analyzed the expression of various proteins related to autophagy and inflammation. We found that the mice treated with FXN had a significantly increased expression of proteins involved in autophagy induction including pCAMKK2 (Ser 511) and pPRKAA2 (Thr 172), and downregulation of pMTOR (Ser 2448). On the similar lines the expression of autophagy initiation and execution proteins pULK1(Ser 317), BECN1, LC3B-II, and ATG5 was found to be significantly increased. The most important impact of autophagy induction by FXN in the hippocampus was the clearance of Aβ_42_. We found only traces of Aβ_42_ in the hippocampus of FXN treated mice in comparison to untreated control. Interestingly, reduced Aβ_42_ levels were directly related to low-reactive phenotype of astrocytes in the hippocampi as evidenced by reduced expression of GFAP. Furthermore, the expression of microglial marker IBA1 was also found to be significantly down regulated in mice treated with FXN. These data indicate that FXN through autophagy not only cleared Aβ_42_ but it also helped mitigation of overall inflammatory microenvironment in the brain. These data were further supported by down regulation of NF-κB and other proteins associated with the activation of NLRP3 inflammasome in the hippocampus. Moreover, these effects of the FXN were not localized to hippocampi as the analysis of proinflammatory cytokines TNF-α, IL-6, and IL-1β in the cortex region of the brain clearly established the strong effect of FXN against neuroinflammation. These data directly correspond to the improvement of memory behavior.

## Conclusion

The analysis of whole data indicates that FXN exerts a potent autophagic effect through activation of PRKAA2 pathway, which helps in the clearance of Aβ_42_ and inhibition of neuroinflammation. In AD, deposition of Aβ_42_ and neuroinflammation together inflict severe damage to neuronal tissue in the brain. Interestingly, FXN exerted these anti-Alzheimer effects at the minimal dose of 5 mg/kg, which corresponds to approximately 25 mg/kg of human equivalent dose, which is a minimum prescribed dose of FXN for OCD. Therefore, FXN through its ability to work against both these key pathological hallmarks of AD at a minimal dose can raise a new hope and warrants its further development against AD.

## Declarations

### Ethics approval and consent to participate

All experiments involving mice were ethically approved by Institutional Animal Ethics Committee (IAEC) of CSIR-IIIM, Jammu, India. Detailed information regarding approval of *in-vivo* and *ex-vivo* experiments is given in Materials and Methods section.

### Consent for publication

Not applicable

### Availability of data and materials

All data generated or analyzed during this study are included in this published article and its supplementary information files.

## List of abbreviations

Aβ: Amyloid-beta
AD: Alzheimer’s disease
PRKAA2: Protein kinase AMP-activated catalytic subunit alpha 2
ATG: Autophagy related
ANOVA: Analysis of variance
ATP: Adenosine triphosphate
ASC: Apoptosis-associated speck-like protein containing a Caspase recruitment domain
BECN1: Beclin 1
BSA: Bovine serum albumin
Baf. A1: Bafilomycin A1
CAMKK: Calcium/calmodulin-dependent protein kinase kinase
CASP1: Caspase-1
NF-κB: Nuclear factor kappa B subunit
DAMPs: Damage-associated molecular patterns
DAPI: 4,6-diaminido-2-phenylindole
DMEM: Dulbecco’s modified eagle medium
DMSO: Dimethyl Sulfoxide
ECL: Enhanced chemiluminescence
EDTA: Ethylenediamine tetra acetic acid
ELISA: Enzyme-linked immunosorbent assay
FBS: Fetal bovine serum
FDA: Federal drug administration
FXN: Fluvoxamine maleate
GFAP: Glial fibrillary acidic protein
HEPES: 4-(2-hydroxyethyl)-1-piperazineethanesulfonic acid
HRP: Horseradish peroxidase
IBA1: Ionized calcium-binding adapter molecule 1
IHC: Immunohistochemistry
LAMP-1: Lysosomal-associated membrane protein 1
LC3: Light chain 3
LPS: Lipopolysaccharide
MTOR: Mammalian target of rapamycin
NFT: Neurofibrillary tangles
NLR: NOD-like receptor
NLRP3: NLR family pyrin domain containing 3
OCD: Obsessive-compulsive Disorder
PBS: Phosphate buffer saline
PEG: Polyethylene glycol
PFA: Paraformaldehyde
PMSF: Phenylmethylsulfonyl fluoride
PVDF: Polyvinylidene difluoride
PYD: Pyrin domain
Rap: Rapamycin
RIPA: Radioimmunoprecipitation assay
SDS: Sodium dodecyl sulfate
SDS-PAGE: SDS poly acrylamide gel electrophoresis
SQSTM1: Sequestosome 1
ULK-1: unc-51-like kinase 1

## Acknowledgements

We are thankful to University Grants Commission (UGC), India for providing the research fellowship to Ms. Sukhleen Kaur.

## Authors’ contributions

SK did most of the in vitro experiments along with some in vivo experiments. KS performed animal behavior studies. KKS and AS performed ELISA experiments. RA and PPS synthesized and characterized fluvoxamine maleate. SMA performed confocal microscope studies. PR helped in genotyping of 5XFAD mice. ZA helped in designing the study. SK and AK designed the study, analyzed the data and wrote the manuscript.

## Disclosure statement

The authors declare that they have no competing interests.

## Funding

The funding for this project was provided by CSIR-Indian Institute of Integrative Medicine (CSIR-IIIM) through the project MLP-6002 (WP-4).

## Notes

### Competing Interest Statement

The authors have declared no competing interest.

### Summary of Updates

The revised version of the manuscript contains a revised Figure 3, which was corrected to replace one western blot in Section B of this Figure. Apart from that figure legends were also aligned correctly in Section A of this Figure.

