## Supplemental Data for "Fluvoxamine maleate ameliorates Alzheimer disease pathology by mitigating amyloid-beta load and neuroinflammation in 5XFAD mice"

### General Scheme for the synthesis of Fluvoxamine maleate

The synthesis of fluvoxamine maleate was done by a well established method comprising of three steps with 5-methoxy-1-(4-(trifluoromethyl)phenyl)pentan-1-one as a starting material (**I**). In the first step compound (**I**) undergoes hydroxylation to form oxime (**II**) as an intermediate. The oxime (**II**) was then alkylated to give fluvoxamine base (**III**) which then undergoes salt formation with maleic acid yielding fluvoxamine maleate (**IV**) as the final desired product [1, 2]. The schematic representation for the synthesis of fluvoxamine maleate (**IV**) is shown below:

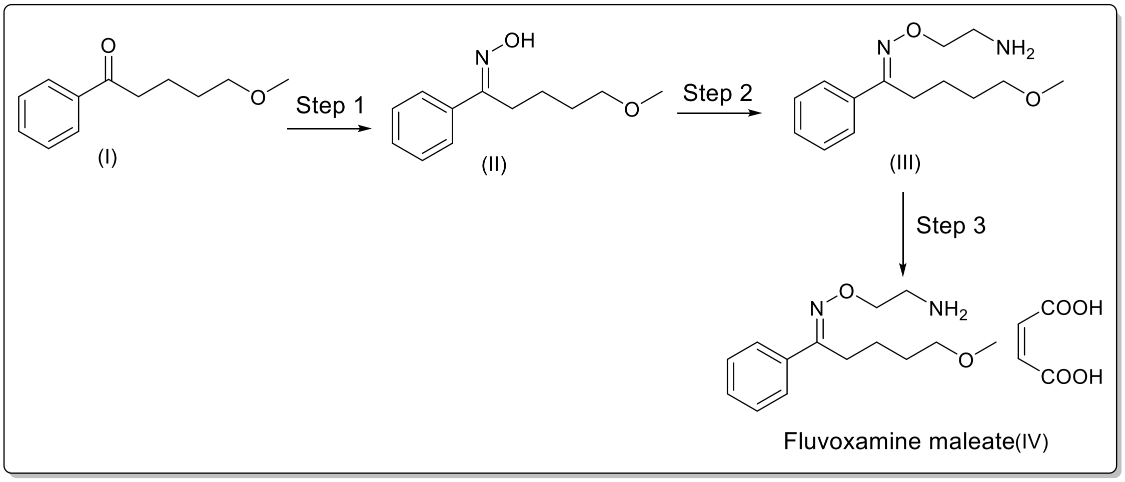


**Scheme 1.** Synthesis of Fluvoxamine maleate.

#### Spectra of Compounds

**$^1$H NMR of Fluvoxamine maleate:** $^1$H NMR (400 MHz, DMSO-$d^6$) δ 7.80 (d, J= 8Hz, 2H), 7.61(d, J= 8Hz, 2H), 6.18(s, 2H) 4.48-4.36 (m, 2H), 3.28-3.22 (m, 11H), 2.81(m, 2H), 1.52(m, 4H); $^{19}$F NMR (400 MHz, DMSO-$d^6$) -62.78.

### $^{1}H$ NMR of Fluvoxamine maleate

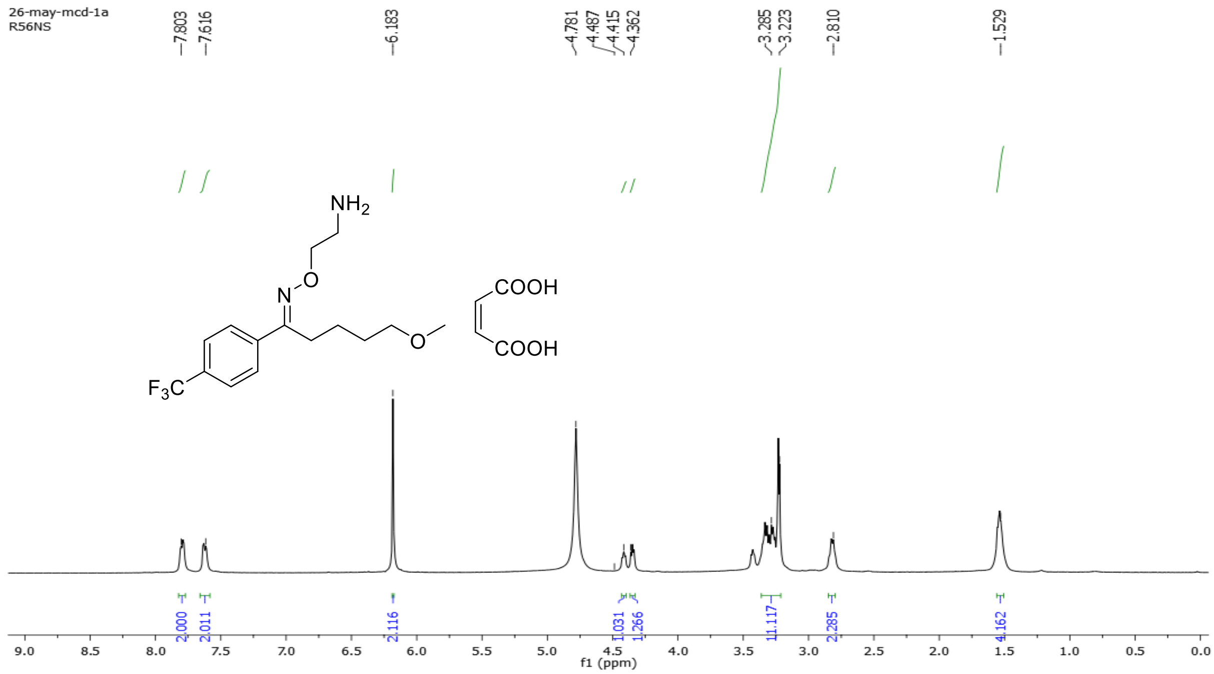


### $^{19}F$ NMR Spectra of Fluvoxamine maleate

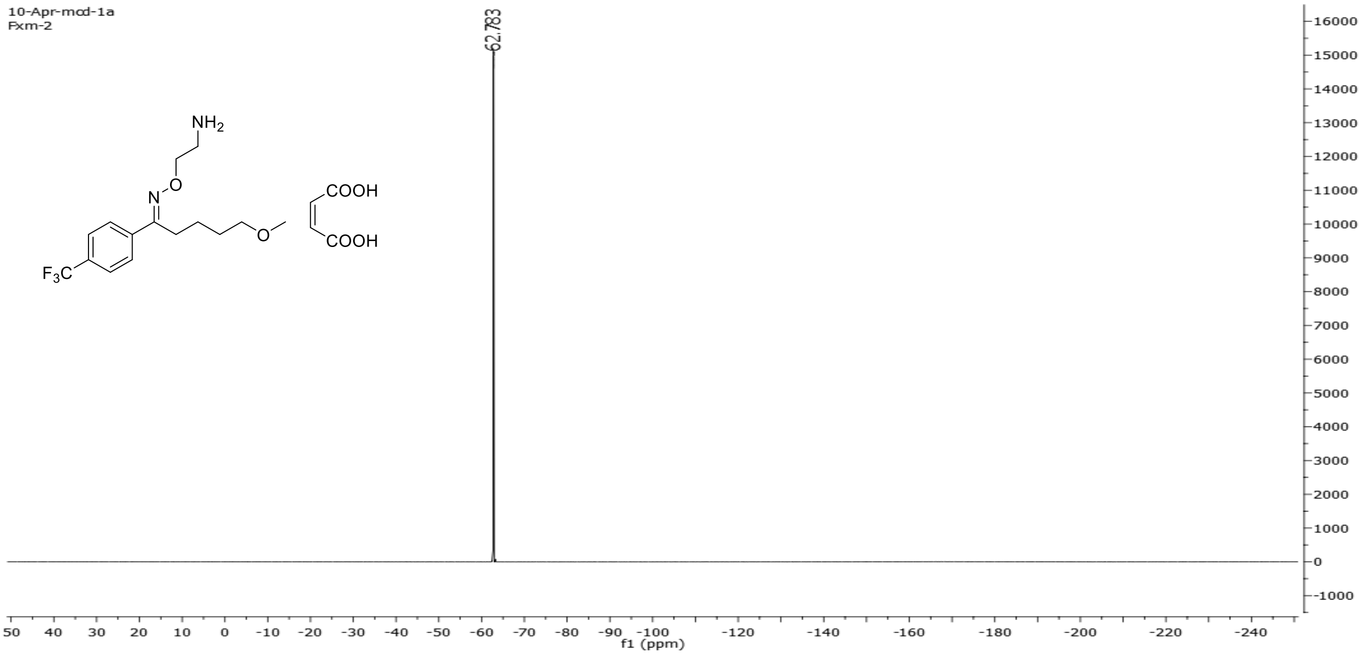

**Figure S1**

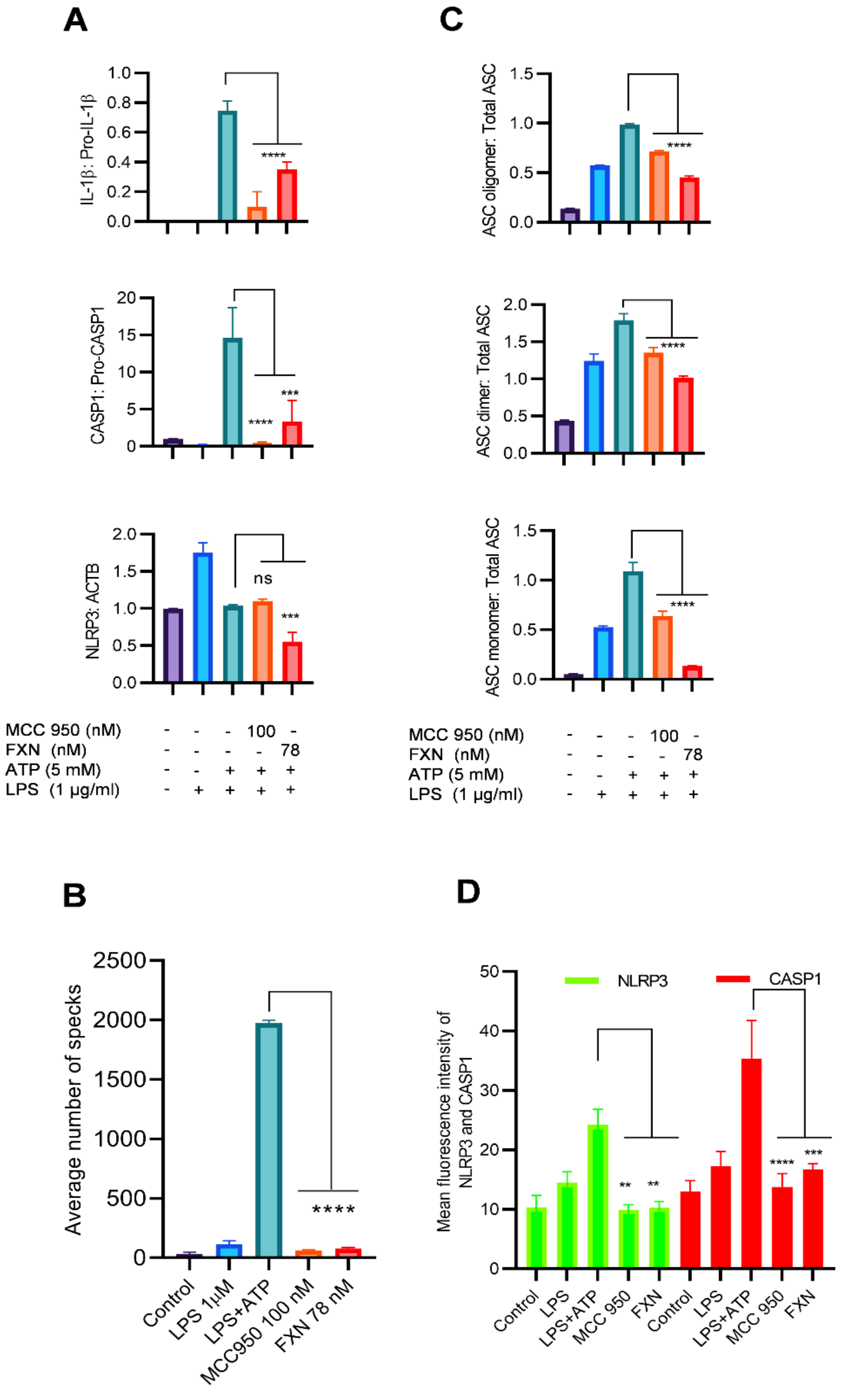


**Figure S1: FXN displayed a potent anti-NLRP3 inflammasome activity in primary astrocytes.**

(A) Densitometric analysis of immunoblots in **Figure 1D**- IL-1β: Pro-IL-1β, CASP1: Pro-CASP1, NLRP3: ACTB**.** (B) Graph representing average number of specks in primary astrocytes shown in **Figure 1E**. (C) Densitometric analysis of immunoblots shown in **Figure 1F**- ASC oligomer, ASC dimer and ASC monomer normalized with total ASC. (D) Graph representing the mean fluorescence intensities of NLRP3 and CASP1 in the images provided in **Figure 1G**. The statistical analysis was performed using one-way ANOVA, followed by post-hoc Bonferroni test. The p-value <0.05 was considered to be statistically significant with values assigned as ****p < 0.0001, ***p < 0.001, **p< 0.01, *p< 0.05 and ns= not significant.

**Figure S2**

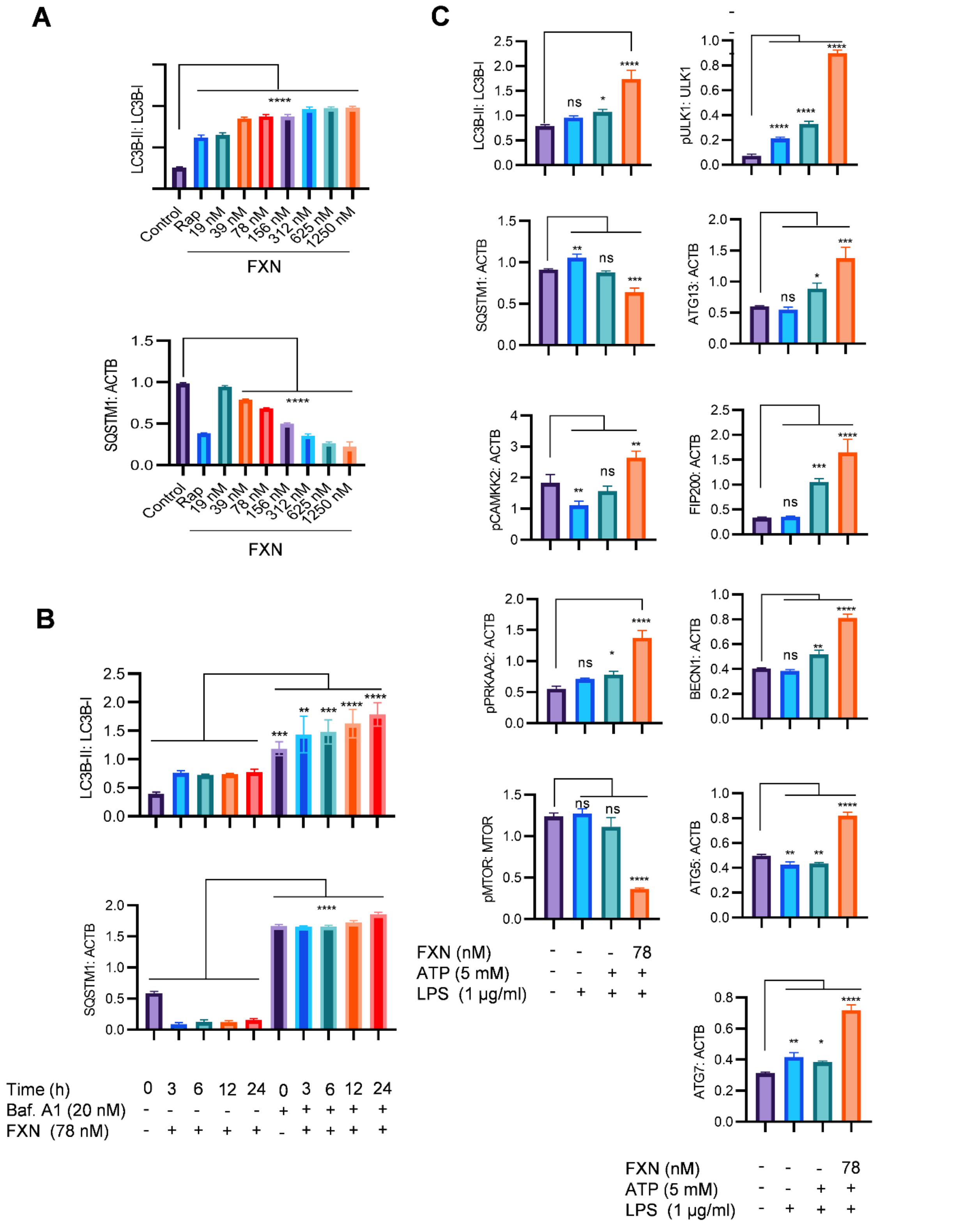


**Figure S2: FXN induced the autophagy under the inflammatory conditions in primary astrocytes.**

Graphs showing Densitometric analysis of immunoblots (A) shown in **Figure 4A**- LC3B-II and SQSTM1 (B) **Figure 4B-** LC3B-II and SQSTM1 (C) **Figure 4D and 4E**- LC3B-II: LC3B-I, SQSTM: ACTB, pCAMKK2: ACTB, pAMPK: ACTB, pMTOR: MTOR, pULK1: ULK1, ATG13: ACTB, FIP200: ACTB, BECN1: ACTB, ATG5: ACTB, and ATG7: ACTB. The statistical analysis was performed using one-way ANOVA, followed by post-hoc Bonferroni test. The p-value <0.05 was considered to be statistically significant with values assigned as ****p < 0.0001, ***p < 0.001, **p< 0.01, *p< 0.05 and ns= not significant.

**Figure S3**

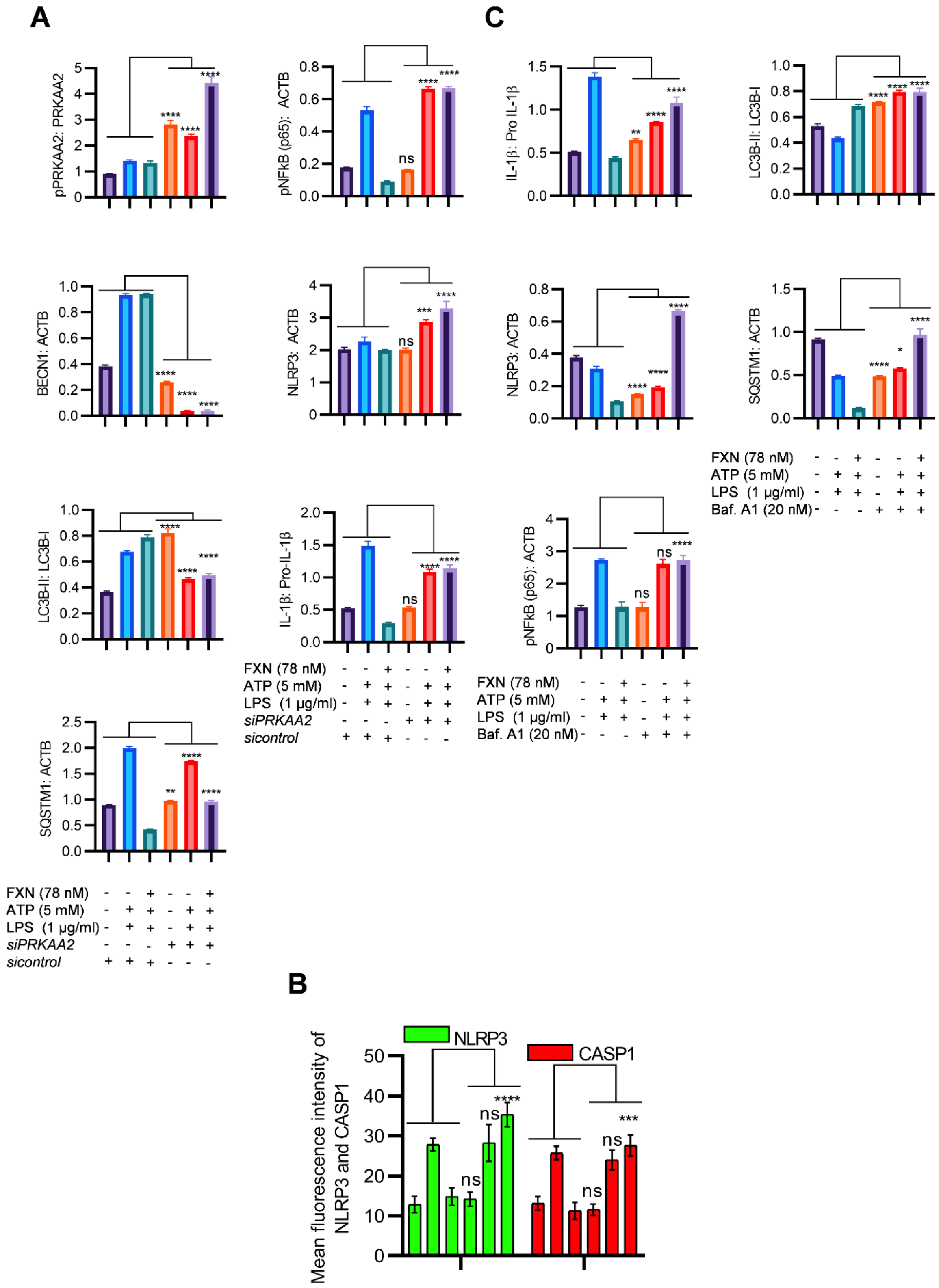


**Figure S3: FXN inhibited NLRP3 inflammasome by inducing autophagy**

(A) Densitometric analysis of immunoblots given in **Figure 5A-** pAMPK: ACTB, BECN1: ACTB, LC3B-II: LC3B-I, SQSTM: ACTB, pNF-κB (p65): ACTB, IL-1β: Pro-IL-1β and NLRP3: ACTB. (B) Graph depicting the mean fluorescence intensities of NLRP3 and CASP1 in the images provided in **Figure 5B**. (C) Densitometry of immunoblots provided in **Figure 5D**- IL-1β: Pro-IL-1β, NLRP3: ACTB, pNF-κB (p65): ACTB, LC3B-II: LC3B-I, SQSTM: ACTB. The statistical analysis was performed using one-way ANOVA, followed by post-hoc Bonferroni test. The p-value <0.05 was considered to be statistically significant with values assigned as ****p < 0.0001, ***p < 0.001, **p< 0.01, *p< 0.05 and ns= not significant.

**Figure S4**

**A**

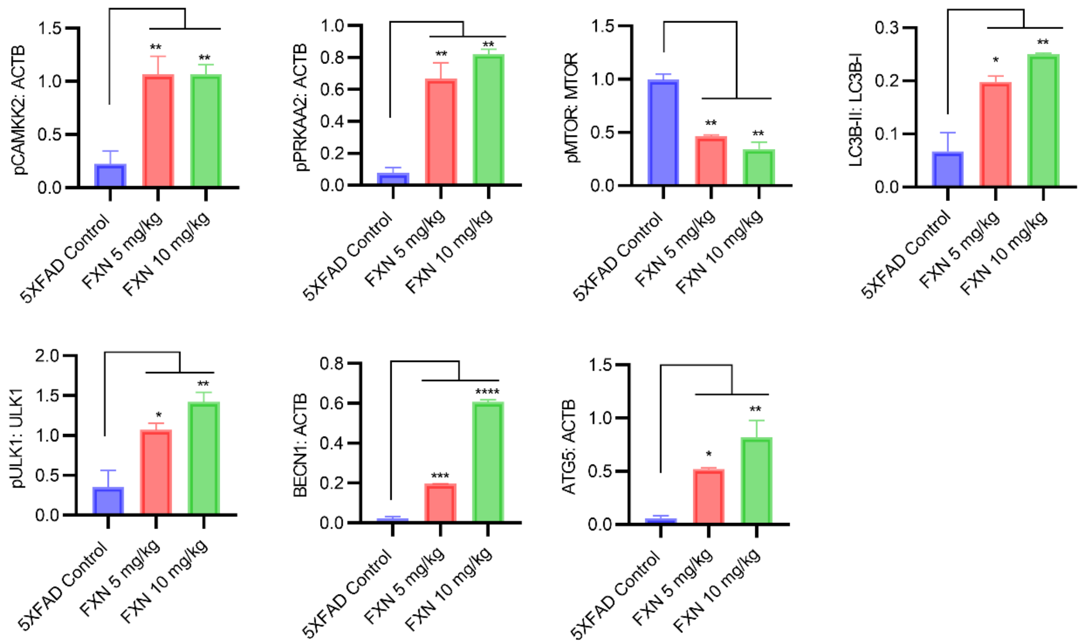


**B**

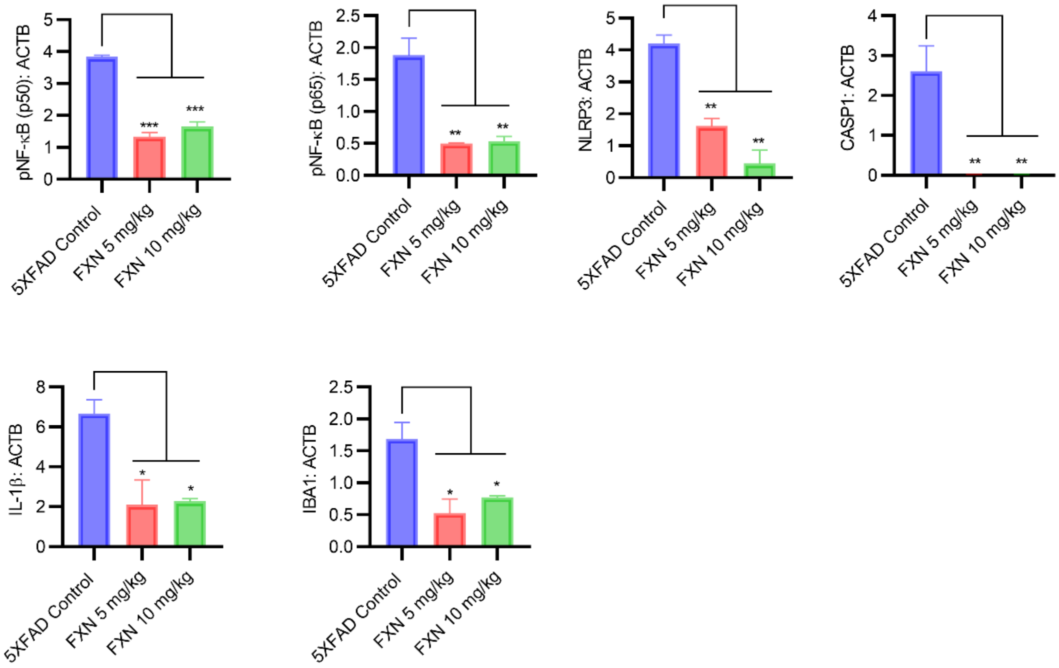


**Figure S4: FXN ameliorated the AD pathology in 5XFAD mice through autophagy-mediated reduction in amyloid beta levels and neuroinflammation.**

Densitometric analysis of immunoblots (A) shown in **Figure 9B-** pCAMKK2: ACTB, pAMPK: ACTB, pMTOR: MTOR, LC3B-II: LC3B1, pULK: ULK1, BECN1: ACTB and ATG5: ACTB (B) **Figure 9C-** NF-κB (p65), NF-κB (p50), NLRP3, CASP1, IL-1β and IBA1 normalized with ACTB. The statistical analysis was performed using one-way ANOVA, followed by post-hoc Bonferroni test. The p-value <0.05 was considered to be statistically significant with values assigned as ****p < 0.0001, ***p < 0.001, **p< 0.01, *p< 0.05 and ns= not significant.
